# Genome-wide gene-environment analyses of major depressive disorder and reported lifetime traumatic experiences in UK Biobank

**DOI:** 10.1101/247353

**Authors:** Jonathan R.I. Coleman, Wouter J. Peyrot, Kirstin L. Purves, Katrina A.S. Davis, Christopher Rayner, Shing Wan Choi, Christopher Hübel, Héléna A. Gaspar, Carol Kan, Sandra Van der Auwera, Mark James Adams, Donald M. Lyall, Karmel W. Choi, Major Depressive Disorder Working Group of the Psychiatric Genomics Consortium, Erin C. Dunn, Evangelos Vassos, Andrea Danese, Barbara Maughan, Hans J. Grabe, Cathryn M. Lewis, Paul F. O’Reilly, Andrew M. McIntosh, Daniel J. Smith, Naomi R. Wray, Matthew Hotopf, Thalia C. Eley, Gerome Breen

## Abstract

Depression is more frequent among individuals exposed to traumatic events. Both trauma exposure and depression are heritable. However, the relationship between these traits, including the role of genetic risk factors, is complex and poorly understood. When modelling trauma exposure as an environmental influence on depression, both gene-environment correlations and gene-environment interactions have been observed. The UK Biobank concurrently assessed Major Depressive Disorder (MDD) and self-reported lifetime exposure to traumatic events in 126,522 genotyped individuals of European ancestry. We contrasted genetic influences on MDD between individuals reporting and not reporting trauma exposure (final sample size range: 24,094-92,957). The SNP-based heritability of MDD was greater in participants reporting trauma exposure (24%) than in individuals not reporting trauma exposure (12%), taking into account the strong, positive genetic correlation observed between MDD and reported trauma exposure. The genetic correlation between MDD and waist circumference was only significant in individuals reporting trauma exposure (r_g_ = 0.24, p = 1.8×10^-7^ versus r_g_ = −0.05, p = 0.39 in individuals not reporting trauma exposure, difference p = 2.3×10^-4^). Our results suggest that the genetic contribution to MDD is greater when additional risk factors are present, and that a complex relationship exists between reported trauma exposure, body composition, and MDD.

## Introduction

Depression is among the most common mental illnesses worldwide and accounts for 5.5% of all years lost through disability globally ^1^. In England approximately 28% of individuals self-report depression during their lifetime ^2^. The most common clinically recognised form of depression is called Major Depressive Disorder (MDD). Both environmental and genetic factors influence MDD. In particular, MDD is more commonly observed among individuals reporting exposure to stressful life events and early-life traumas ^3–6^. In turn, reported trauma exposure has been robustly correlated with a range of adverse life outcomes including MDD ^6–9^. The relationship between MDD and reported trauma exposure is complex, with studies showing both that reported trauma exposure is associated with subsequent MDD, and that MDD is associated with subsequent reported trauma exposure ^10, 11^. However, the majority of people reporting exposure to traumatic experiences do not report MDD ^6–9^.

Twin studies show that MDD is moderately heritable, with 30-40% of the variance in MDD attributable to genetic factors ^12^. The proportion of heritability captured by common genetic variants, also known as single nucleotide polymorphism or SNP-based heritability, can be estimated from genome-wide association study (GWAS) data. Such estimates tend to be lower than those obtained from twin approaches, due to the incomplete capture of genetic information in GWAS data among other reasons ^13^. The most recent major depression GWAS from the Psychiatric Genomics Consortium was anchored in 35 cohorts (including the 23andMe discovery cohort ^14^) recruited with a variety of methods ^15^. This meta-analysis identified 44 loci significantly associated with major depression, and estimated a SNP-based heritability of 9-10% ^15^. GWAS results strongly suggest both the mild and more severe forms of depression are polygenic, with potentially thousands of variants with very small individual effects contributing to risk.

There are far fewer genetic studies of reported trauma exposure than of MDD. However, the available studies have demonstrated that reported trauma exposure is heritable, with twin heritability estimates of 20-50% ^16–18^ and SNP-based heritability estimates of 30% ^19^. Combining measures of trauma exposure and depression on a large scale is difficult and, as with many environmental measures, requires careful phenotyping ^20^. Potential confounds include the (often unavoidable) use of retrospective self-reported measures of trauma exposure, which can be weakly correlated with objective measures of traumatic experiences ^9^. Furthermore, current (i.e. state) low mood or MDD can increase self-reporting of previous trauma exposure ^9, 21^. Previous individual study cohorts have generally been too small for effective GWAS, while meta-analyses have contained considerable heterogeneity due to the use of different phenotyping instruments in the included studies.

However, some notable genome-wide analyses of MDD and trauma exposure have been performed. A genome-wide by environment interaction study of depressive symptoms and stressful life events in 7,179 African American women identified a genome-wide association near the *CEP350* gene (although this did not replicate in a smaller cohort) ^22^. A recent investigation in 9,599 Han Chinese women with severe MDD identified three variants associated with MDD specifically in individuals who did not report exposure to traumatic events prior to MDD onset ^23^.

Several attempts have been made to estimate the interaction of overall genetic risk and trauma by using polygenic risk scores for MDD to perform polygenic risk score-by-trauma interaction analyses. Such studies test whether there are departures from additivity (where the combined effect of risk score and trauma differs from the sum of the individual effects) or from multiplicativity (where the combined effect differs from the product of the individual effects). Results from these have been highly variable, with reports of significant additive and multiplicative interactions ^24^; significant multiplicative interactions only ^25^; and, in the largest previous study published (a meta-analysis of 5,765 individuals), no interactions ^26^.

Studies of gene-environment interaction usually assume the genetic and environmental influences are independent and uncorrelated ^27^. However, genetic correlations between reported trauma exposure and MDD have been reported, both from twin studies ^28–30^ and from the genomic literature ^22, 26^. Reports of the magnitude of this genetic correlation have varied widely, which reflects differences in defining trauma exposure, and in the populations studied. While some studies have identified a very high genetic correlation (95%) ^22^, others have found no such correlation ^23^. The relationship between reported trauma exposure and MDD, and the influence of genetic variation on that relationship, is therefore complex and unresolved.

The release of mental health questionnaire data from the UK Biobank resource provides an opportunity to assess the relationship between genetic variation, risk for MDD, and reported trauma exposure in a single large cohort. We performed GWAS of probable MDD with and without reported lifetime trauma exposure in UK Biobank European ancestry individuals. These GWAS were underpowered to detect individual genetic loci at the very low effect sizes seen in previous GWAS of MDD. However, these results enabled us to estimate the SNP-based heritability of MDD in individuals with and without reported lifetime trauma exposure, and so to test whether the relative contribution of genetic influences to MDD differs in the context of reported trauma exposure. We then estimated the genetic correlation between MDD in the two groups, to assess whether the genetic component differed between individuals reporting and not reporting trauma exposure. In addition, we compared the patterns of genetic correlation between MDD in these groups and a wide range of physical and psychiatric traits, in order to assess whether the genetic relationship of MDD to other traits varies in the context of reported trauma exposure. Finally, we performed polygenic risk scoring, using external traits commonly comorbid with MDD, and sought to extend previous analyses of PRS-by-trauma interactions in MDD.

## Methods

### Phenotype definitions

The UK Biobank assessed a range of health-related phenotypes and biological measures including genome-wide genotype data in approximately 500,000 British individuals aged between 40 and 70 ^31^. This includes 157,366 participants who completed an online follow-up questionnaire assessing common mental health disorders, including MDD symptoms, and 16 items assessing traumatic events (Resource 22 on http://biobank.ctsu.ox.ac.uk) ^32^. Phenotypes were derived from this questionnaire, using definitions from a recent publication describing its phenotypic structure ^32^.

Individuals with probable MDD met lifetime criteria based on their responses to questions derived from the Composite International Diagnostic Interview (CIDI; Supplementary Table 1). We excluded cases if they self-reported previous diagnoses of schizophrenia, other psychoses, or bipolar disorder. Controls were excluded if they self-reported any mental illness, reported taking any drug with an antidepressant indication, had previously been hospitalised with a mood disorder or met previously-defined criteria for a mood disorder (Supplementary Table 1) ^33^.

Participants were asked questions relating to traumatic experiences in childhood using the Childhood Trauma Screener (a shortened version of the Childhood Trauma Questionnaire ^34–36)^ and an equivalent screener for adulthood developed by the UK Biobank Mental Health steering group to mirror the childhood items ^32^. In addition, participants were asked questions related to events that commonly trigger post-traumatic stress-disorder (PTSD). Responses to individual questions (items) in these three categories (child trauma, adult trauma, PTSD-relevant trauma) were dichotomised and compared between MDD cases and controls (Supplementary Table 2a).

We selected reported items with an odds ratio > 2.5 with MDD, to obtain a single binary variable for stratification that captured exposure to the traumatic events most associated with MDD. Items from all three trauma categories were reported more in MDD cases compared to controls. Of the selected items, three referred to events in childhood (did not feel loved, felt hated by a family member, sexually abused). Another three items referred to events in adulthood (physical violence, belittlement, sexual interference), and one item assessed a PTSD-relevant event (ever a victim of sexual assault). In order to capture increased severity of exposure, only individuals reporting two or more of these items were included as reporting trauma exposure. Individuals reporting none of the items were included as not reporting trauma exposure. Individuals reporting a single trauma item, or who did not provide an answer were excluded from the analyses (Supplementary Table 1). A breakdown of reported traumatic experiences by sex and MDD status is provided in Supplementary Table 2b. Further discussion of the definition of trauma exposure is included in the Supplementary Note.

### Phenotype preparation for analyses

Three sets of analyses comparing MDD cases and controls were performed (i) overall, (ii) limited to individuals reporting trauma exposure, and (iii) limited to individuals not reporting trauma exposure (Table 1). In addition, sensitivity analyses were performed on reported trauma exposure (overall and stratified by MDD diagnosis; see Supplementary Methods and Results, and Supplementary Table 3). For each analysis, phenotypes were first residualised on 6 ancestry principal components from the genetic data of the European samples as well as factors capturing initial assessment centre and genotyping batch. More details on phenotype preparation can be found in the Supplementary Methods.

**Table 1:** Participants available for analysis. Groups of individuals used in each of the three analyses are in bold. The superscripts denote the groups used in each of the three main analyses: a) MDD in all participants (29,475 cases, 63,482 controls, N = 92,957); b) MDD in participants reporting trauma exposure (13,393 cases, 10,701 controls, N = 24,094); c) MDD in participants not reporting trauma exposure (9,487 cases, 39,677 controls, N = 49,164).

|  |  | Participants with genomic data |  |  |  |
| --- | --- | --- | --- | --- | --- |
|  |  | Reported trauma exposure | No reported trauma exposure | Excluded | Total |
| MDD | Cases | **13,393<sup>b</sup>** | **9,487<sup>c</sup>** | 6,595 | **29,475<sup>a</sup>** |
|  | Controls | **10,701<sup>b</sup>** | **39,677<sup>c</sup>** | 13,104 | **63,482<sup>a</sup>** |

### Phenotype distribution

Previous analyses have shown that, compared to the participants in the UK Biobank as a whole, those who completed the mental health questionnaire were more likely to have a university degree, came from a higher socioeconomic background, and reported fewer long-standing illnesses or disabilities ^32^. Accordingly, participants were compared across a number of standard demographic variables and common correlates of MDD: sex, age (at questionnaire), education (university degree vs. not), neighbourhood socioeconomic status (SES, as Townsend deprivation index ^37^) and BMI (recorded from measurements taken at the initial recruitment of the participants into the biobank). For further details on these analyses, see Supplementary Methods.

### Genetic data

Genetic data for GWAS analyses came from the full release of the UK Biobank data (N=487,410; ^38^). Autosomal genotype data from two highly-overlapping custom genotyping arrays (covering ∼800,000 markers) underwent centralised quality control before being imputed in a two-stage imputation to the Haplotype Reference Consortium (HRC) and UK10K (for rarer variants not present in the HRC) reference panels ^38–40^. In addition to this central quality control, variants for analysis were limited to common variants (minor allele frequency > 0.01) that were either directly genotyped or imputed from the HRC with high confidence (IMPUTE INFO metric > 0.4) ^39^.

Individuals were excluded where recommended by the UK Biobank core analysis team for unusual levels of missingness or heterozygosity, or if they had withdrawn consent for analysis. Using the genotyped SNPs, individuals with call rate < 98%, who were related to another individual in the dataset (KING r < 0.044, equivalent to removing up third-degree relatives and closer ^41^) or whose phenotypic and genotypic gender information was discordant (X-chromosome homozygosity (F_X_) < 0.9 for phenotypic males, F_X_ > 0.5 for phenotypic females) were also excluded. Removal of relatives was performed using a “greedy” algorithm, which minimises exclusions (for example, by excluding the child in a mother-father-child trio). All analyses were limited to individuals of European ancestry, as defined by 4-means clustering on the first two genetic principal components provided by the UK Biobank ^42^. This ancestry group included 95% of the respondents to the mental health questionnaire - as such, the non-European ancestry groups were considered too small to analyse informatively. Principal components analysis was also performed on the European-only subset of the data using the software flashpca2 ^43^. After quality control, individuals with high-quality genotype data and who had completed the online mental health questionnaire were retained for analysis (N=126,522).

GWAS analyses used the imputed data as described above. Genetic correlation analyses used the results of the GWAS analyses. Polygenic risk score analyses and SNP-based heritability analyses in BOLT-LMM used the genotyped variants ^38^. These latter analyses were limited to common variants (minor allele frequency > 0.01) with call rate >98% that were in approximate Hardy-Weinberg equilibrium (HWE test p > 10^-8^). The same individuals were used for analyses using the imputed and the genotyped data.

### Analyses

#### Genome Wide Association Studies (GWAS)

GWAS were performed to assess the association of individual genetic variants with MDD. These analyses were first undertaken for the entire sample regardless of reported trauma exposure, then stratified by reported trauma exposure. GWAS were performed using linear regressions on imputed genotype dosages in BGenie v1.2 ^38^, with residualised phenotypes as described above. Phenotypes and genotypes were mean-centred and standardised. Genome-wide significance was defined at the conventional level p < 5 x 10^-8 44^. Results from each GWAS were clumped to define genetic loci in PLINK2 ^45^. Loci were defined following established protocols (Supplementary Methods) ^15^.

Betas from the GWAS were converted to odds ratios (OR) using LMOR (http://cnsgenomics.com/shiny/LMOR/) and observed sample prevalences ^46^. Standard errors were calculated from the p-value and estimated OR ^47^. Performing GWAS on residuals, rather than including covariates in the analysis, is a restriction imposed by the BGenie software (which was used because it is specifically designed for analysing the UK Biobank genetic data). Sensitivity analyses were performed to test for biases resulting from this method. Specifically, for each GWAS, each variant with nominal significance (p<0.0001) was also tested using logistic regression including covariates in R 3.4.1, in order to confirm the results from BGenie ^48^.

#### SNP-based heritability

Results from GWAS were combined to assess the proportion of variance due to the additive effect of common genetic variants (SNP-based heritability). SNP-based heritability was calculated on the observed scale using BOLT-LMM v2.3 ^49^. The estimate for MDD in the cohort was converted to the liability scale in R 3.4.1, assuming a population prevalence of 28% ^2, 50^. Converting estimates of SNP-based heritability for a case-control trait from the observed scale to the liability scale requires accurate estimates of the lifetime prevalence of the trait in the (sub)population. When comparing a trait stratified by a correlated variable (as is the case when we compare the SNP-based heritability of MDD stratified by reported trauma exposure), the population prevalence in each stratum is unknown. To address this, we approximated the expected prevalence of MDD in individuals either reporting or not reporting trauma exposure (Supplementary Methods). This allowed us to convert the observed scale SNP-based heritability of MDD to the liability scale in both strata (i.e. those reporting and those not reporting trauma exposure). A second challenge is that trauma exposure is itself a heritable trait that is genetically correlated with MDD in this study. The potential impact of this on SNP-based heritability estimation is not intuitive. To benchmark our findings, we performed simulations of SNP-level data to explore the expected SNP-based heritability of MDD in individuals reporting and not reporting trauma exposure, assuming differences in SNP-based heritability resulted only from the genetic correlation between MDD and reported trauma exposure. Further details of these analyses are provided in the Supplementary Methods.

#### Genetic correlations

Genetic correlations (r_g_) were calculated to assess shared genetic influences between MDD and other phenotypes, using GWAS summary statistics and LD Score regression v1.0.0 ^51^ using the default HapMap LD reference. Two sets of genetic correlations were calculated. First, we calculated genetic correlations between the phenotypes examined within this paper (internal phenotypes). We calculated the genetic correlation between MDD and reported trauma exposure in the full dataset, and then the genetic correlation between MDD in individuals reporting trauma exposure and MDD in individuals not reporting trauma exposure. Secondly, we also calculated genetic correlations between each GWAS from this analysis and a curated list of 308 publicly-available phenotypes (external phenotypes) ^51, 52^.

Genetic correlations were tested for difference from 0 (default in LD Score), and for difference from 1 (in Microsoft Excel, converting r_g_ to a chi-square as [(r_g_- 1)/se]^2^) ^51, 52^. Genetic correlations were considered significant if they passed the Bonferroni-adjusted threshold for the effective number of traits studied in each analysis (internal: p < 0.01; external: p < 2.5×10^-4^). The effective number of traits was calculated as the number of principal components explaining 99.5% of the variance in the pairwise genetic correlation matrix (internal: 5; external: 202). External phenotype GWAS all had heritability estimates such that h^2^/SE > 2, and produced valid (i.e. non-NA) r_g_ with all other phenotypes tested.

The genetic correlation of MDD with each external phenotype was compared between individuals reporting trauma exposure and individuals not reporting trauma exposure using a two-stage method. First, differences were assessed using two sample z-tests ^53^. Nominally-significant differences (p < 0.05) by this method were then compared using the block-jackknife (Supplementary Methods) ^52, 54, 55^. Results using the jackknife were considered significant if they passed the Bonferroni-adjusted threshold (p < 2.5×10^-4^).

#### Polygenic Risk Scoring

Polygenic risk scores were calculated to further assess shared genetic influences between MDD and traits known to be correlated to MDD. Specifically, risk scores from analyses of major depression (MDD) ^15^, schizophrenia (SCZ) ^56^, bipolar disorder (BIP) ^57^, body mass index (BMI) ^58^ and glycated haemoglobin (HbA1c; used as a negative control) ^59^ were calculated and compared in all participants and stratifying by reported trauma exposure. The PGC major depression GWAS contained participants from UK Biobank, so to derive the MDD risk score we used a restricted set of summary statistics without these individuals (but including individuals from 23andMe, whose diagnoses were self-reported ^14^). For further discussion of this overlap, see Supplementary Note ^15^. Risk scores were calculated using PRSice v2 (https://github.com/choishingwan/PRSice ^45, 60^) at seven thresholds (external GWAS p < 0.001, 0.05, 0.1, 0.2, 0.3, 0.4 and 0.5) to allow assessment of the spread of association between risk score and MDD. Analyses used logistic regression, including all covariates used in creating the residuals for GWAS. In total, five external phenotypes were used to produce risk scores for the three target phenotypes (MDD overall, and stratified by reported trauma exposure/non-exposure), resulting in 15 analyses. A conservative Bonferroni adjustment for multiple testing was used, correcting for 105 tests (seven thresholds and 15 analyses), giving a final threshold for significance of p < 0.0004.

In addition to these stratified analyses, we performed formal risk score-by-environment analyses to estimate the effect on MDD of the interaction between genetic variants across the whole genome (modelled as a risk score) and reported trauma exposure. We calculated interactions between reported trauma exposure and the risk score capturing the most variance from each of the main-effect polygenic risk score analyses. These analyses included the same covariates used in the GWAS, and all risk score-by-covariate and reported trauma exposure-by-covariate interactions ^61, 62^. Both multiplicative and additive interactions were tested. A significant multiplicative interaction means that the combined effect of the risk score and reported trauma exposure differs from the product of their individual effects. Multiplicative interactions were tested using logistic regression ^25, 26^. A significant additive interaction means that the combined effect of the risk score and reported trauma exposure differs from the sum of their individual effects. Additive interactions were tested using linear regression (Supplementary Methods).

### Sensitivity analyses

Differences in phenotypic variables were observed between cases and controls. To assess the impact of including these variables as covariates, all analyses were rerun retaining all previous covariates and including as further covariates: age (at questionnaire), neighbourhood socioeconomic status (SES, as Townsend deprivation index ^37^), BMI (at baseline assessment), and a binary variable of education (university degree vs. not). The same covariates were also included in polygenic risk score and SNP-based heritability analyses. Sensitivity analyses focussing on reported trauma exposure as an outcome were similarly rerun (Supplementary Methods).

The majority of the sample with data on both MDD symptoms and reported trauma status were controls who did not report trauma (Table 1). To assess whether this disbalance in sample status affected our results, genetic correlation analyses with external phenotypes were rerun on ten downsampled cohorts, each with 9,487 participants (the number of cases not reporting trauma exposure; see Supplementary Methods).

In order to test whether our definition of trauma exposure affected the main finding of our paper, we performed three further sensitivity analyses, redefining reported trauma exposure. First, we assessed if our main finding was robust to changing the threshold for including MDD-relevant trauma, by redefining reported trauma exposure as a report of i) one or more and ii) three or more of the seven MDD-relevant trauma items. Second, we assessed whether the timing of trauma exposure affected this finding by redefining reported trauma exposure as a report of iii) one or more of the five childhood trauma items. We then re-analysed the heritability of MDD in individuals reporting and not reporting trauma exposure using these three alternative definitions.

## Results

### Phenotype distribution

Phenotypic and genetic data were available on 24,094 to 92,957 individuals (Table 1). Overall, 36% of individuals met our definition of MDD-relevant trauma exposure, and were more frequently cases (45%) than controls (17%; OR = 5.23; p < 10^-50^, chi-square test). We assessed a number of phenotypic correlates of depression to confirm that these correlates differed between MDD cases and controls, and to assess whether these differences were affected by trauma exposure. Cases differed significantly from controls overall. Individuals with MDD were mostly females, significantly younger, less likely to have a university degree, came from more deprived neighbourhoods, and had higher BMI at recruitment. These differences persisted when the cohort was limited just to individuals reporting trauma exposure, and when the cohort was limited just to individuals not reporting trauma exposure. Furthermore, cases reporting trauma exposure differed from cases not reporting trauma exposure, in that they were mostly females, younger, more likely to have a degree (note difference from case-control comparisons), came from more deprived neighbourhoods, and had higher BMI at recruitment. The same differences (in the same direction) were observed between controls reporting and not reporting trauma exposure (all p < 0.05; Supplementary Table 4).

### Genome-wide association studies

We performed GWAS for MDD overall and stratified by reported trauma exposure to obtain results for heritability and genetic correlation analyses (Supplementary Table 6; Supplementary Figures 1-3). No analysis showed evidence of genome-wide inflation that was attributable to confounding (95% confidence intervals of all regression intercepts from LD Score included 1; Supplementary Table 7). One genome-wide significant locus (rs11515172, Chr 9:11Mb, p = 3.82×10^-8^) was identified in the analysis of MDD overall, and remained significant when using logistic regression (p = 4.69 x 10^-8^, OR = 0.96, SE = 0.007; Supplementary Table 6). This locus has been repeatedly associated with depression ^15, 63, 64^ and with neuroticism ^65–68^; however, it should be noted that all of these studies included UK Biobank. The locus is intergenic, and is not annotated to any currently known biological feature of interest (Supplementary Table 8).

**Figure 1:**
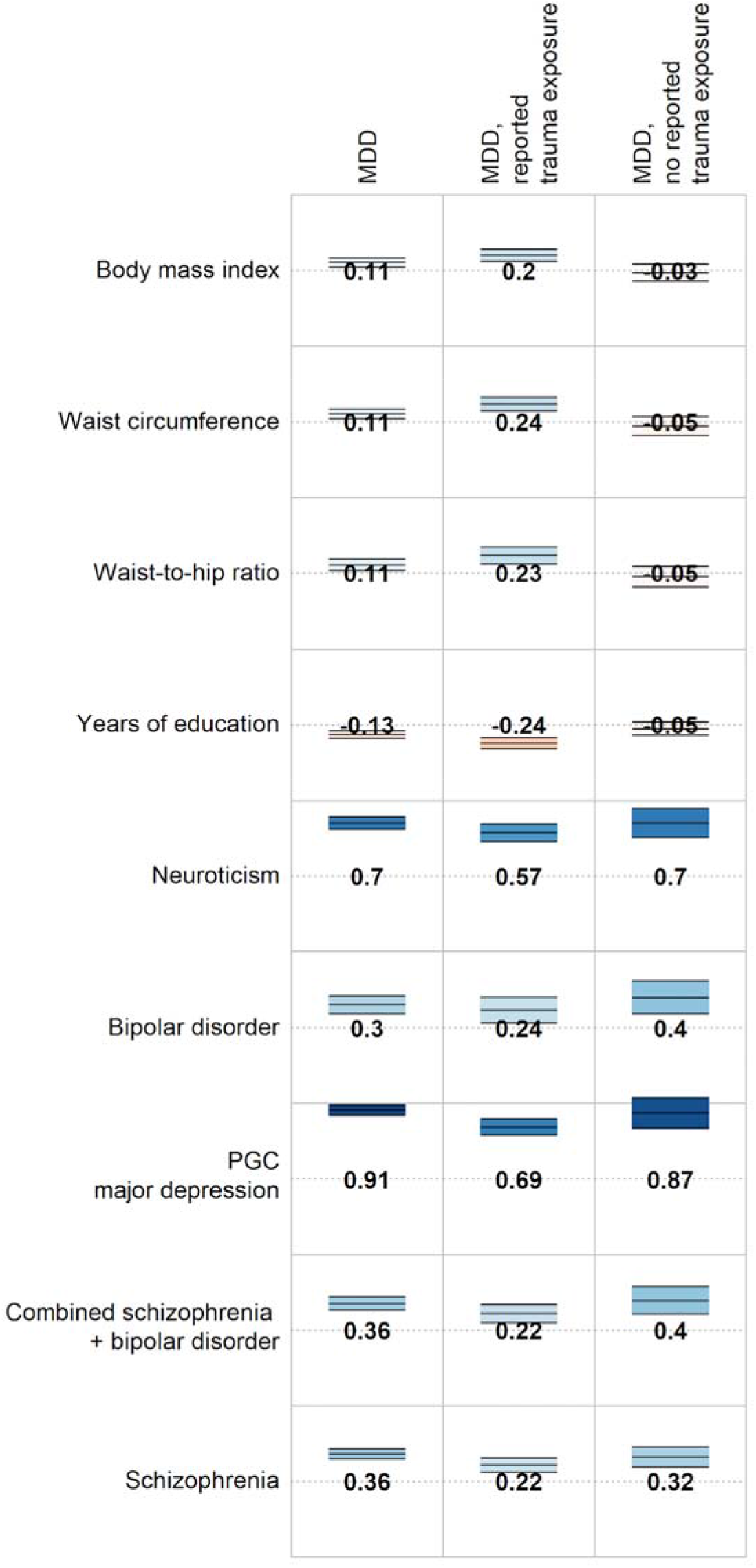
Genetic correlations between MDD (overall and stratified by reported trauma exposure) and selected traits and disorders. Full genetic correlation results are available in Supplementary Table 11. Numbers = genetic correlations. Colour = direction of effect (blue = positive, red = negative). Colour intensity = size of correlation. Upper and lower bars are 95% confidence interval of genetic correlation.

### SNP-based heritability

First we estimated the observed scale SNP-based heritability of MDD overall and stratified by reported trauma exposure. Second, in order to assess whether the relative influence of genetic variants on MDD differed by reported trauma status, we converted SNP-heritabilities to the liability scale. We assumed a prevalence of 28% for self-reported MDD in the full population ^2^. Based on this, and on the ratio of MDD cases:controls in the sample, we estimated the prevalence of MDD in the trauma-exposed population as 52%, and in the unexposed population as 17%. Using these estimates of population prevalence, the liability scale estimate of MDD SNP-based heritability was 20% (95% confidence interval: [18-22%]) overall. In those reporting trauma exposure, the liability scale SNP-based heritability of MDD was 24% [18-31%], and in those not reporting trauma exposure it was 12% [7-16%]. The SNP-based heritability of MDD was significantly greater in individuals who reported trauma exposure compared to those who did not (p = 0.0021, Z-test).

These estimated SNP-heritabilities could be confounded by genetic correlation between MDD and reported trauma exposure. We conducted simulations of SNP-level data to quantify the expected difference in SNP-based heritability from genetic correlation alone. Our simulations yielded expected estimates for the liability scale SNP-based heritability of MDD of 14-15% in those reporting trauma exposure, and 15-16% in those not reporting trauma exposure (Supplementary Methods). This small difference in expected SNP-based heritability for those reporting and not reporting trauma is in the opposite direction to our findings. This suggests that our findings cannot be explained by genetic correlation between MDD and reported trauma exposure, nor by the transformation from the observed scale to the liability scale.

### Genetic correlations

Genetic correlations were calculated between MDD and reported trauma to explore the genetic relationship between these traits. Further genetic correlations were calculated between MDD in the two strata to assess whether genetics influences on MDD differ in the context of reported trauma exposure (Supplementary Table 10).

We observed a significant r_g_ between MDD and reported trauma exposure in the full cohort (0.62 [95% CI: 0.76-0.94], p < 10^-50^). Given that trauma items were selected for association with MDD, we also calculated the genetic correlation between MDD in the full cohort and reported trauma exposure in just the controls, which was also significantly greater than 0 (0.31 [0.18-0.45], p = 4×10^-6^; Supplementary Table 10). This correlation persisted when using independent major depression GWAS summary statistics, as reported trauma exposure was significantly correlated with the MDD polygenic risk score (Spearman’s rho = 0.0675, p < 10^-50^) ^15^. The genetic correlation between MDD in individuals reporting trauma exposure and MDD in individuals not reporting trauma exposure was high and did not differ significantly from 1 (r_g_ = 0.77 [0.48-1.05]; difference from 0: p = 1.8 x 10^-7^; difference from 1: p = 0.11).

Genetic correlations were calculated between MDD and all available external traits to systematically assess whether genetic relationships with MDD differed in the context of reported trauma exposure. All psychiatric traits included were significantly associated (p < 2.5×10^-4^) with MDD, but this association did not differ substantially in magnitude between the groups reporting and not reporting trauma exposure (z-test for comparisons of r_g_ - _Δ_r_g_ - ranged from p = 0.10 - 0.99; Figure 1). In contrast, waist circumference was significantly associated with MDD only in individuals reporting trauma exposure (r_g_ = 0.24), and the correlation was significantly larger than that in individuals not reporting trauma exposure (r_g_ = −0.05, jackknife p_Δrg_ = 2.3×10^-4^). Other correlations between MDD and body composition, reproductive, and socioeconomic phenotypes were larger in the group reporting trauma exposure compared to individuals not reporting trauma exposure, but these differences did not remain significant following multiple testing correction (all jackknife p > 2.5×10^-4^; Figure 1, Supplementary Table 11).

### Polygenic risk scores across strata

We performed polygenic risk score analyses to further explore how stratification by trauma status affects the genetic relationship between MDD and specific correlates of MDD, and to mirror previous analyses in the literature (Figure 2, Table 2; see Supplementary Table 12 for full details of all risk score analyses, including the number of SNPs in each score) ^26^. Individuals with high genetic risk scores for MDD were more likely to be cases than controls, and a significant additive interaction term was observed from linear regression. Specifically, the combined effect of the MDD risk score and reported trauma exposure on MDD was greater than the sum of the individual effects (beta > 0, Table 2 central panel). However, the multiplicative interaction term was not significant (p > 0.01). The presence of an interaction on the additive scale reflects the greater SNP-based heritability of MDD in individuals reporting trauma exposure (SNP-h^2^ = 24%) compared to those not reporting trauma exposure (SNP-h^2^ = 12%), as described above.

**Figure 2:**
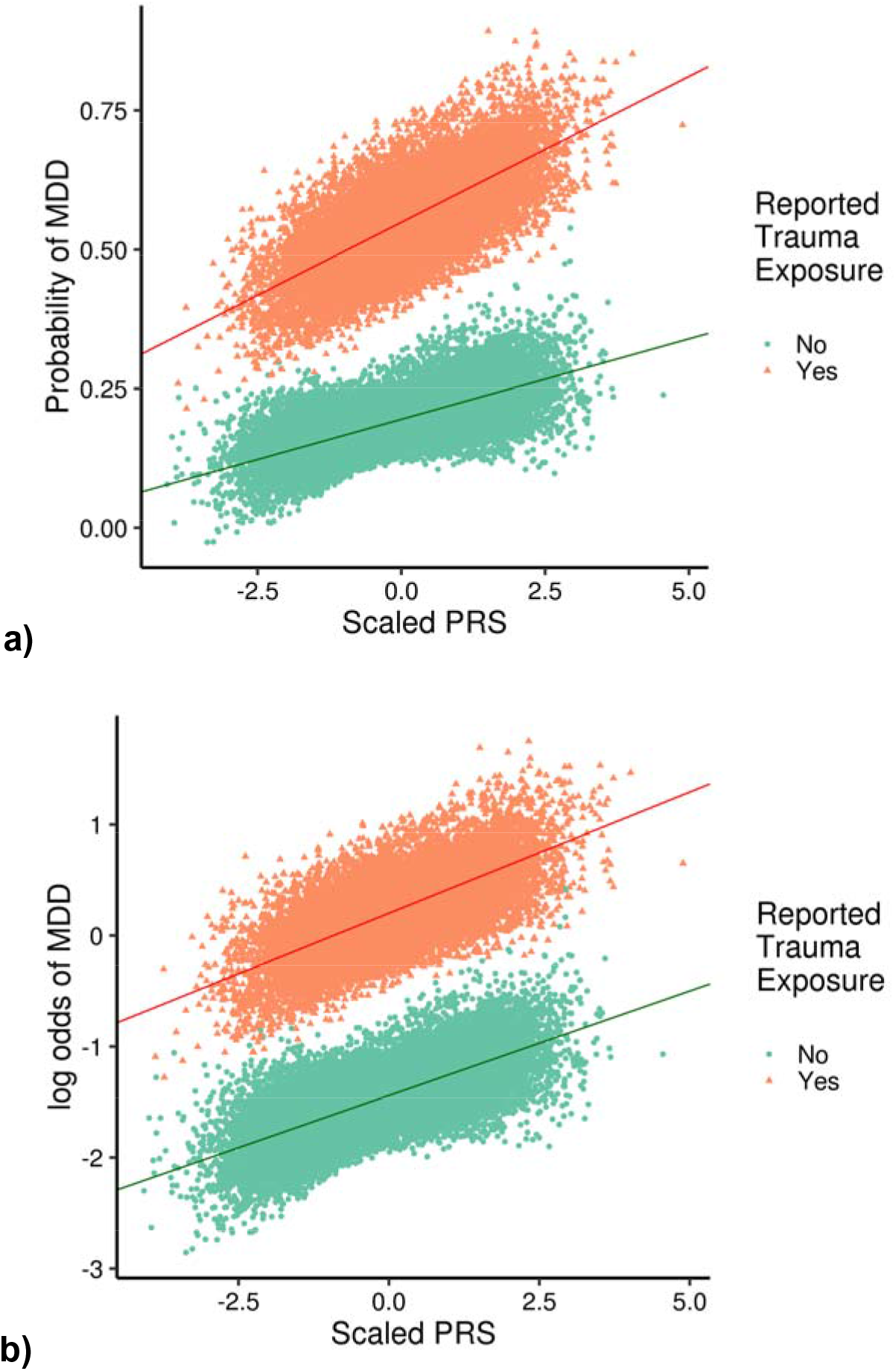
Association between MDD polygenic risk score (PRS) and MDD. Individuals reporting trauma exposure are shown as orange triangles, and those not reporting trauma exposure as green dots. Panel a shows the relationship on the linear additive scale, and panel b shows the relationship on the multiplicative scale. A significant interaction is observed on the additive scale only, as shown by differing slopes of the two regression lines in panel a.

**Table 2:** Main effect and interaction effects for polygenic risk scores (PRS) associated with MDD overall and in stratified analyses. Interaction effects are on he additive scale (Beta) and the multiplicative scale (OR). Bold = significant associations (main analyses: p < 0.000143; interactions: p < 0.01). Base N = ases // Controls. OR/Beta = Increase with 1 SD increase in risk score or trauma exposure. Results are reported at the “best” threshold (that with the lowest p-value in main effect analyses) - results across all thresholds are reported in Supplementary Table 12.

| Base | Base N | Best Threshold | Analysis | PRS |  |  | PRS x Reported Trauma |  |  |  |  |  |
| --- | --- | --- | --- | --- | --- | --- | --- | --- | --- | --- | --- | --- |
|  |  |  |  | OR | 95% CI | p | Additive |  |  | Multiplicative |  |  |
|  |  |  |  |  |  |  | Beta | 95% CI | p | OR | 95% CI | p |
| MDD | 116,404 // 314,990 | **0.5** | *MDD* | **1.26** | **1.24-1.28** | **< 10<sup>-50</sup>** | **0.011** | **0.008-0.014** | **2.69x10<sup>-11</sup>** | 1.01 | 1.00 - 1.03 | 0.132 |
| SCZ | 36,989 // 113,075 | **0.3** |  | **1.11** | **1.09-1.12** | **1.11 x 10<sup>-41</sup>** | 0.008 | -0.003-0.004 | 0.659 | 0.99 | 0.97-1.00 | 0.158 |
| BIP | 7,481 // 9,250 | **0.2** |  | **1.07** | **1.05-1.08** | **4.57 x 10<sup>-19</sup>** | -0.000 | -0.003-0.003 | 0.961 | 0.99 | 0.97-1.00 | 0.165 |
| BMI | 339,224 | **0.3** |  | **1.04** | **1.03-1.06** | **2.60 x 10<sup>-8</sup>** | **0.006** | **0.003-0.009** | **1.13x10<sup>-4</sup>** | **1.02** | **1.01-1.04** | **0.0074** |
| HB1Ac | 46,368 | 0.001 |  | 1.01 | 1.00-1.02 | 0.163 | 0.002 | -0.001-0.005 | 0.186 | 1.01 | 0.99-1.02 | 0.391 |
| MDD | 116,404 // 314,990 | **0.4** | *MDD, reported trauma exposure* | **1.24** | **1.21-1.27** | **< 10<sup>-50</sup>** |  |  |  |  |  |  |
| SCZ | 36,989 // 113,075 | **0.5** |  | **1.06** | **1.03-1.09** | **2.96 x 10<sup>-5</sup>** |  |  |  |  |  |  |
| BIP | 7,481 // 9,250 | 0.5 |  | 1.04 | 1.01-1.07 | 0.00329 |  |  |  |  |  |  |
| BMI | 339,224 | **0.5** |  | **1.07** | **1.04-1.10** | **1.85 x 10<sup>-7</sup>** |  |  |  |  |  |  |
| HB1Ac | 46,368 | 0.001 |  | 1.02 | 1.00-1.05 | 0.0863 |  |  |  |  |  |  |
| MDD | 116,404 // 314,990 | **0.4** | *MDD, no reported trauma exposure* | **1.21** | **1.18-1.23** | **< 10<sup>-50</sup>** |  |  |  |  |  |  |
| SCZ | 36,989 // 113,075 | **0.5** |  | **1.09** | **1.06-1.11** | **4.34 x 10<sup>-12</sup>** |  |  |  |  |  |  |
| BIP | 7,481 // 9,250 | **0.2** |  | **1.07** | **1.04-1.09** | **4.05 x 10<sup>-8</sup>** |  |  |  |  |  |  |
| BMI | 339,224 | 0.3 |  | 1.02 | 1.00-1.04 | 0.0980 |  |  |  |  |  |  |
| HB1Ac | 46,368 | 0.001 |  | 1.01 | 0.99-1.03 | 0.492 |  |  |  |  |  |  |

In contrast, although those with higher BMI risk scores were more likely to be cases than controls, this only passed correction for multiple testing in individuals reporting trauma exposure. Both the additive (beta > 0) and the multiplicative (OR > interaction terms were significant, suggesting the combined effect on MDD from BMI risk score and reported trauma exposure together was greater than expected from both the sum of the individual risks and from their product, respectively (OR > 1).

Individuals with high genetic risk scores for SCZ were more likely to be cases than controls, but this did not differ between strata (both interaction terms p >0.01). Individuals with higher BIP risk scores were also more likely to be cases than controls - although this association was not significant in the subset of individuals reporting trauma exposure, no significant interaction term was observed, suggesting the observed difference in results within-strata may be due to differences in power. No significant differences were observed in the negative control analysis with HbA1c.

### Sensitivity analyses

Four sets of sensitivity analyses were performed. In the first set, all analyses were repeated using reported trauma exposure as the phenotype, assessed overall and stratified by MDD (as opposed to the primary analysis, where MDD was the phenotype and analyses were stratified by reported trauma exposure). Results from these analyses were broadly similar to the results from the primary analysis (Supplementary Tables 3-12, Supplementary Figures 4-7).

The second set of sensitivity analyses repeated the primary analyses with additional covariates to assess the impact of controlling for age, neighbourhood socioeconomic status, BMI, and education. This did not alter the conclusions drawn from the GWAS and SNP-based heritability analyses, nor from the genetic correlations observed between the internal phenotypes (those assessed in this study; Supplementary Tables 13-18). Genetic correlations between MDD and external phenotypes did not differ significantly from the main analysis (all z-test p < 0.05), but were sufficiently attenuated that the genetic correlations of MDD with waist circumference was no longer significantly different between individuals reporting and not reporting trauma exposure. Differences in the polygenic risk score analyses were limited to analyses involving the BMI risk score. The BMI risk score was no longer associated with MDD in any analysis, and no interactions including the BMI risk score remained significant. This suggests the effects of the BMI risk score in the main analysis likely result from BMI differences between MDD cases and controls, rather than being true effects on MDD.

The third set of sensitivity analyses repeated the genetic correlation analyses, but downsampled the analysed cohort such that each of the four groups (MDD cases/controls reporting/not reporting trauma exposure) had 9,487 participants (the size of the smallest group from the main analysis, cases not reporting trauma exposure). In these analyses, genetic correlations between MDD and external phenotypes were attenuated across most phenotypes, but not significantly (two-sample z-tests, all p > 0.05; Supplementary Table 19). As such, the general pattern of genetic correlations observed in the main analysis was retained, although the genetic correlations of MDD with waist circumference was no longer significantly different between individuals reporting and not reporting trauma exposure.

The final set of sensitivity analyses repeated the SNP-based heritability analyses of MDD in individuals reporting and not reporting trauma exposure, altering the definition of reported trauma exposure in three ways (increasing and decreasing the number of items required to be defined as reporting trauma exposure, and limiting the items considered to only childhood experiences). The purpose of these analyses was to test the robustness of our key finding (greater MDD SNP-based heritability in trauma-exposed individuals compared to those not reporting trauma exposure). Neither increasing nor decreasing the number of MDD-relevant items selected, nor focussing on childhood items, altered our conclusions (Supplementary Table 20).

Full results for all four sensitivity analyses are included in the Supplementary Material.

## Discussion

We investigated the relationship between MDD and self-reported trauma exposure in the largest single cohort available to date (N=73,258 with MDD and reported trauma data). We examined individual genetic variants, SNP-based heritability, genetic correlations, and polygenic risk scores. The SNP-based heritability of MDD was higher in individuals reporting trauma exposure than in individuals not reporting trauma exposure. This was not attributable to gene-environment correlation, nor to the transformation of SNP-based heritability from the observed to the liability scale. Despite the significant difference in SNP-based heritability across the two strata, the genetic correlation between MDD in individuals reporting and not reporting trauma exposure was not statistically different from 1. Polygenic risk score-by-reported trauma exposure interaction analyses identified significant interactions for both MDD and BMI risk scores. However, the interactions involving the BMI risk score appear to reflect differences in measured BMI between MDD cases and controls, rather than differences directly related to MDD. Finally, a significant genetic correlation between MDD and waist circumference was observed only in individuals reporting trauma exposure, and was absent from those not reporting trauma exposure.

A number of limitations should be considered when assessing our results. Our simulations suggest that the significantly higher SNP-based heritability of MDD in individuals reporting trauma exposure did not result from gene-environment correlation between MDD and reported trauma exposure, nor from the transformation of the observed scale SNP-based heritability to the liability scale. However, we could not address further sources of potential bias. These could arise from genetic architectures other than those simulated (including non-additive genetic effects), from intrinsic challenges of heritability estimation in case-control data (including ascertainment bias and the effects of covariates not included in the model) ^69, 70^, or from potential collider bias (the induction of a correlation between two uncorrelated variables by conditioning on a third unknown variable correlated with both) resulting from selection bias ^71^. We also assumed that the population prevalence of reported trauma exposure can be extrapolated from that observed in this sample (see Supplementary Methods). Although the UK Biobank allows us to integrate genetic and environmental data at scale, and is a reasonably homogeneous cohort, it also has a “healthy volunteer bias”, whereby the participants tend to have better overall health and higher socioeconomic status compared to the equivalent overall population of this age ^72^. It is possible that the depressive and traumatic experiences reported by these participants may not generalise to the whole population, or to clinically-ascertained cases. Furthermore, we focussed on European ancestry; further studies in non-European populations are required ^73^.

To obtain further insight into the association of genome-wide genetic variation and reported trauma exposure with MDD (and to enable comparison with previous studies ^24–26)^, we carried out polygenic risk score-by-environment interaction analyses. There are a number of limitations to consider when interpreting such analyses. Polygenic risk score-by-environment interaction analyses test a specific hypothesis, namely that the overall association of common variants with the outcome (modelled as a risk score) varies dependent on the environmental exposure being tested. As such, the absence of an interaction does not preclude the existence of specific variant-by-environment interactions, including those featuring variants contributing to the risk score. Furthermore, we cannot exclude the possibility that the correlation between the MDD risk score and reported trauma exposure may alter the observed interaction. As such, caution is needed before drawing strong conclusions, especially given the small effect sizes, and limited predictive power, of risk scores in this study (Supplementary Table 12).

Throughout this paper, we have referred to our depression phenotype as “MDD” rather than “major depression”. We do this because our definition is based on the CIDI-SF, which has previously been shown to have good concordance with direct clinical assessments of MDD ^74, 75^. However, it should be noted that direct assessment was not performed, and our probable MDD cases may not have met criteria within a clinical setting. Nonetheless, genetic correlations between studies of MDD and of major depression are high, suggesting there is strong genetic continuity across different methods of assessing depression ^15, 64^.

Trauma exposure was defined in this study using retrospective self-report. This is not the ideal measure for this phenotype, and precludes robust measurement of the severity and timing of the reported trauma exposure. However, retrospective report is the only feasible option in large cohorts like the UK Biobank. The requirement for cohort sizes large enough to identify the small individual genetic effects typical of complex genetic traits such as MDD makes self-report the most practicable method of data-collection. Retrospectively reported trauma and MDD data are not robust to reverse causation, and our results cannot strongly inform any temporal or causal hypotheses about the relationship between trauma and MDD. Such hypotheses could be tested using longitudinal studies (with the inherent logistical difficulties in obtaining both environmental and genomic data) or through more powerful genomic studies of trauma exposure in a larger cohort. This latter design would enable the identification of sufficient robustly associated genetic variants to inform approaches such as Mendelian randomisation (which we were underpowered to examine in this study). In addition, future work may benefit from assessing the heritability of broader depression phenotypes that lie beyond our binary criteria, including reward sensitivity and negative valence traits ^76^.

Our findings suggest that the genetic variants associated with MDD are the same in individuals reporting and not reporting trauma exposure, because the genetic correlation between MDD measured in these two groups was not significantly different from 1. However, the SNP-based heritability of MDD was *greater* in individuals reporting compared to not reporting trauma exposure. This suggests that the combined effect of the variants associated with MDD is greater in people reporting trauma exposure than in those who do not. The mechanism underlying this finding is uncertain. One possibility is that exposure to traumatic events might amplify genetic influences on MDD beyond the magnitude of the effects seen in the absence of trauma (consistent with the stress-diathesis hypothesis ^77–79)^. The concept that genetic variance varies with exposure to different environments is well-recognised in studies of animal populations in the wild ^80^. However, the opposite may also be true; genetic influences on MDD could increase an individual’s likelihood of experiencing and/or reporting of trauma, and through doing so increase the apparent heritability of MDD by partly incorporating genetic influences related to trauma reporting itself ^11^. A final possibility relates to the components of variance involved in calculating SNP-based heritability. Phenotypic variance can be attributed either to the SNPs measured in the GWAS, or to environmental sources of variance reflecting all phenotypic variance not explained by common variants. It is possible that the genetic component of variance is constant across the strata, but that the environmental component is smaller in individuals reporting trauma exposure, for example due to the shared (and thus more similar/less variable) exposure of these individuals to MDD-relevant traumatic experiences. This would result in greater heritability in individuals reporting trauma exposure. We provide these explanations as potential interpretations of these findings. However, these are not the only possibilities, and it is also likely that multiple such mechanisms are involved.

In polygenic risk score-by-reported trauma exposure interaction analyses, we identified a significant interaction on the additive scale for the combined effect of the MDD risk score and reported trauma exposure on risk of MDD. These results are also reflected in the larger SNP-based heritability of MDD in exposed compared to unexposed individuals. The simplest explanation for this result is that the effects of the MDD risk score and reported trauma exposure on MDD combine multiplicatively, such that their combined effects are greater than the sum of their individual effects. For the BMI risk score however, the interaction with reported trauma exposure appears to be more complex, combining neither additively nor multiplicatively. In sensitivity analyses controlling for BMI (obtained at recruitment, approximately five years before the mental health questionnaire), the BMI risk score-by-reported trauma exposure interaction was no longer significant, suggesting that the observed interaction can be explained by differences in measured BMI. Further research, with concurrent measurements of BMI, trauma exposure and MDD in a longitudinally-sampled cohort would offer further insight into the relationship between these three variables.

The high genetic correlation between MDD in individuals reporting and not reporting trauma exposure was supported by significant genetic correlations between MDD and other psychiatric disorders regardless of reported trauma exposure. In individuals reporting trauma exposure, a further significant genetic correlation was observed between MDD and waist circumference, which was significantly greater than the equivalent correlation in those not reporting trauma exposure. Although not significant, there was also a general pattern of higher genetic correlations between MDD and several weight-related measures and educational attainment, in individuals reporting trauma exposure. This is consistent with previous literature on traumatic experiences across the life course, and related phenomena such as Adverse Childhood Experiences (ACEs). This literature has found that such adversities are associated not only with psychiatric risk but also with wide-ranging impairments in social and health outcomes including obesity and education ^81–84^. However, we stress that causal conclusions cannot be drawn from these data, or that the reported trauma exposure is responsible for the observed differences.

Our estimate of the SNP-based heritability of MDD (20%) is higher than that reported in previous studies of major depression (∼9%) ^15^. This may be explained by the relative homogeneity of the UK Biobank compared to previous meta-analyses. The UK Biobank is a single-country cohort ascertained using a consistent protocol. The same questionnaire was used to gather symptom data, and the samples were stored, extracted, and genotyped using a single method. In contrast, meta-analyses have needed to combine diverse ascertainment, sampling, and genotyping; SNP-based heritability has been reported to decrease with increasing numbers of meta-analysed samples ^85^. Previous analyses have assessed alternative depression phenotypes in the UK Biobank ^64^. Our MDD phenotype (based on DSM criteria for MDD) is most similar to the probable MDD phenotype from Howard et al, rather than the less strictly-defined “broad depression” phenotype, which includes those who seek treatment for depression, anxiety and related phenotypes. More specifically the “broad depression” phenotypes includes many people with MDD, but also many people who may not meet diagnostic criteria for MDD or instead have anxiety disorders or a similar closely related common mental disorder. Our MDD phenotype includes only those people who meet diagnostic criteria (using the CIDI questionnaire). Our estimate from BOLT-LMM (19.9%) is within the bounds of the reported range of GREML estimates by geographic region reported for the probable MDD (0% to 27.5%) phenotype. Our LDSC-based estimate is higher than the equivalent from Howard et al (4-5%). However, the estimate from Howard et al is considerably lower than previous estimates ^15^, and potentially lacks both the specificity of definitions from clinical practice or structured questionnaires, and the sensitivity of broad phenotyping methods.

Our results also differ in several respects from those of a study of MDD and adversity in Han Chinese women ^23^. No difference in the SNP-based heritability of MDD between individuals reporting and not reporting trauma exposure was observed in the previous study, and we did not replicate individual variant results. However, this is unsurprising, as there are a number of differences between the studies of which the primary one is sample size (this study: 73,258; CONVERGE: 9,599). Other differences included culture and ethnicity, and the deeper phenotyping methodology applied in CONVERGE, resulting in a severe inpatient MDD phenotype. Notably, the previous study did not report a genetic correlation between MDD and trauma exposure ^23^.

Sensitivity analyses focussed on trauma found that self-reported traumatic experience was significantly heritable, as has been previously observed ^19^. We strongly emphasise that this does not necessarily imply that traumatic experiences themselves have a biological component - such experiences may be associated with other significantly heritable traits, and their biology would then be reflected in the observed heritability of trauma exposure. One potential set of heritable traits that may be associated with reporting traumatic experiences are personality traits such as risk-taking, and this might explain the observed genetic correlations with psychiatric traits. A similar phenomenon has been proposed to underlie observed genetic correlations with socioeconomic status ^86^. Our trauma exposure measure relies on retrospective self-report, which is itself correlated with personality traits and mood at time of report ^9^. This may also explain the genetic correlations we observe with reported trauma exposure (including in controls, who do not report previous psychiatric illness).

In summary, we find that genetic associations with MDD in UK Biobank vary by context. Specifically, the SNP-based heritability of MDD is larger in individuals reporting trauma exposure compared to those not doing so. Furthermore, the genetic correlation of MDD with waist circumference was significant only in individuals reporting exposure to trauma. Nonetheless, a strong genetic correlation was observed between MDD measured in the two strata. Together, these findings suggest the relative contribution of genetic variants to variance in MDD is greater when additional risk factors are present.

## Acknowledgements

We thank the members of the UK Biobank Mental Health Genetics Group for their valuable discussion and feedback on this work. We are also deeply indebted to the scientists involved in the construction of the UK Biobank, and to the investigators who comprise the PGC. Finally, we thank the hundreds of thousands of subjects who have shared their life experiences with investigators in the UK Biobank and the PGC.

This research has been conducted using the UK Biobank Resource, as an approved extension to application 16577 (Dr Breen). This study represents independent research funded by the National Institute for Health Research (NIHR) Biomedical Research Centre at South London and Maudsley NHS Foundation Trust and King’s College London. The views expressed are those of the authors and not necessarily those of the NHS, the NIHR or the Department of Health and Social Care. High performance computing facilities were funded with capital equipment grants from the GSTT Charity (TR130505) and Maudsley Charity (980). WJP was funded by NWO Veni grant 91619152. K.L.P acknowledges funding from the Alexander von Humboldt Foundation. N.R.W acknowledges funding from the Australian National Health and Medical Research Council (1078901 and 1087889). The PGC has received major funding from the US National Institute of Mental Health and the US National Institute of Drug Abuse (U01 MH109528 and U01 MH1095320).

## Notes

#### Summary of Updates

- Inclusion of additional sensitivity analyses increasing/decreasing the threshold for defining reported trauma exposure, and focussing on traumas experienced in childhood. No qualitative alterations to overall result. - Alteration to multiple-testing correction for genetic correlation analyses to better account for correlated phenotypes - Expanded discussion of potential mechanisms of observed increase in MDD SNP-based heritability when trauma exposure is reported - Substantial rewrite to increase clarity

