## Supplementary Information for "Genome-wide gene-environment analyses of major depressive disorder and reported lifetime traumatic experiences in UK Biobank"

Manuel Mattheisen* 6, 7, 8

Maciej Trzaskowski 1

Enda M Byrne 1

Abdel Abdellaoui 9

Mark J Adams 10

Esben Agerbo 11, 12, 13

Tracy M Air 14

Till F M Andlauer 15, 16

Silviu-Alin Bacanu 17

Marie Bækvad-Hansen 13, 18

Aartjan T F Beekman 19

Tim B Bigdeli 17, 20

Elisabeth B Binder 15, 21

Julien Bryois 22

Henriette N Buttenschøn 13, 23, 24

Jonas Bybjerg-Grauholm 13, 18

Na Cai 25, 26

Enrique Castelao 27

Jane Hvarregaard Christensen 8, 13, 24

Toni-Kim Clarke 10

Jonathan R I Coleman 28

Lucía Colodro-Conde 29

Baptiste Couvy-Duchesne 2, 30

Nick Craddock 31

Gregory E Crawford 32, 33

Gail Davies 34

Ian J Deary 34

Franziska Degenhardt 35

Eske M Derks 29

Nese Direk 36, 37

Conor V Dolan 9

Erin C Dunn 38, 39, 40

Thalia C Eley 28

Valentina Escott-Price 41

Farnush Farhadi Hassan Kiadeh 42

Hilary K Finucane 43, 44

Jerome C Foo 45

Andreas J Forstner 35, 46, 47, 48

Josef Frank 45

Héléna A Gaspar 28

Michael Gill 49

Fernando S Goes 50

Scott D Gordon 29

Jakob Grove 8, 13, 24, 51

Lynsey S Hall 10, 52

Christine Søholm Hansen 13, 18

Thomas F Hansen 53, 54, 55

Stefan Herms 35, 47

Ian B Hickie 56

Per Hoffmann 35, 47

Georg Homuth 57

Carsten Horn 58

Jouke-Jan Hottenga 9

David M Hougaard 13,18

David M Howard 10, 28

Marcus Ising 59

Rick Jansen 19

Ian Jones 60

Lisa A Jones 61

Eric Jorgenson 62

James A Knowles 63

Isaac S Kohane 64, 65, 66

Julia Kraft 4

Warren W. Kretzschmar 67

Zoltán Kutalik 68, 69

Yihan Li 67

Penelope A Lind 29

Donald J MacIntyre 70, 71

Dean F MacKinnon 50

Robert M Maier 2

Wolfgang Maier 72

Jonathan Marchini 73

Hamdi Mbarek 9

Patrick McGrath 74

Peter McGuffin 28

Sarah E Medland 29

Divya Mehta 2, 75

Christel M Middeldorp 9, 76, 77

Evelin Mihailov 78

Yuri Milaneschi 19

Lili Milani 78

Francis M Mondimore 50

Grant W Montgomery 1

Sara Mostafavi 79, 80

Niamh Mullins 28

Matthias Nauck 81, 82

Bernard Ng 80

Michel G Nivard 9

Dale R Nyholt 83

Paul F O'Reilly 28

Hogni Oskarsson 84

Michael J Owen 60

Jodie N Painter 29

Carsten Bøcker Pedersen 11, 12, 13

Marianne Giørtz Pedersen 11, 12, 13

Roseann E Peterson 17, 85

Erik Pettersson 22

Wouter J Peyrot 19

Giorgio Pistis 27

Danielle Posthuma 86, 87

Jorge A Quiroz 88

Per Qvist 8, 13, 24

John P Rice 89

Brien P. Riley 17

Margarita Rivera 28, 90

Saira Saeed Mirza 36

Robert Schoevers 91

Eva C Schulte 92, 93

Ling Shen 62

Jianxin Shi 94

Stanley I Shyn 95

Engilbert Sigurdsson 96

Grant C B Sinnamon 97

Johannes H Smit 19

Daniel J Smith 98

Hreinn Stefansson 99

Stacy Steinberg 99

Fabian Streit 45

Jana Strohmaier 45

Katherine E Tansey 100

Henning Teismann 101

Alexander Teumer 102

Wesley Thompson 13, 54, 103, 104

Pippa A Thomson 105

Thorgeir E Thorgeirsson 99

Matthew Traylor 106

Jens Treutlein 45

Vassily Trubetskoy 4

Andrés G Uitterlinden 107

Daniel Umbricht 108

Sandra Van der Auwera 109

Albert M van Hemert 110

Alexander Viktorin 22

Peter M Visscher 1, 2

Yunpeng Wang 13, 54, 104

Bradley T. Webb 111

Shantel Marie Weinsheimer 13, 54

Jürgen Wellmann 101

Gonneke Willemsen 9

Stephanie H Witt 45

Yang Wu 1

Hualin S Xi 112

Jian Yang 2, 113

Futao Zhang 1

Volker Arolt 114

Bernhard T Baune 115, 116, 117

Klaus Berger 101

Dorret I Boomsma 9

Sven Cichon 35, 47, 118, 119

Udo Dannlowski 114

EJC de Geus 9, 120

J Raymond DePaulo 50

Enrico Domenici 121

Katharina Domschke 122, 123

Tõnu Esko 5, 78

Hans J Grabe 109

Steven P Hamilton 124

Caroline Hayward 125

Andrew C Heath 89

Kenneth S Kendler 17

Stefan Kloiber 59, 126, 127

Glyn Lewis 128

Qingqin S Li 129

Susanne Lucae 59

Pamela AF Madden 89

Patrik K Magnusson 22

Nicholas G Martin 29

Andrew M McIntosh 10, 34

Andres Metspalu 78, 130

Ole Mors 13, 131

Preben Bo Mortensen 11, 12, 13, 24

Bertram Müller-Myhsok 15, 132, 133

Merete Nordentoft 13, 134

Markus M Nöthen 35

Michael C O'Donovan 60

Sara A Paciga 135

Nancy L Pedersen 22

Brenda WJH Penninx 19

Roy H Perlis 38, 136

David J Porteous 105

James B Potash 137

Martin Preisig 27

Marcella Rietschel 45

Catherine Schaefer 62

Thomas G Schulze 45, 93, 138, 139, 140

Jordan W Smoller 38, 39, 40

Kari Stefansson 99, 141

Henning Tiemeier 36, 142, 143

Rudolf Uher 144

Henry Völzke 102

Myrna M Weissman 74, 145

Thomas Werge 13, 54, 146

Cathryn M Lewis* 28, 147

Douglas F Levinson* 148

Gerome Breen* 28, 149

Anders D Børglum* 8, 13, 24

Patrick F Sullivan* 22, 150, 151

1, Institute for Molecular Bioscience, The University of Queensland, Brisbane, QLD, AU

2, Queensland Brain Institute, The University of Queensland, Brisbane, QLD, AU

3, Analytic and Translational Genetics Unit, Massachusetts General Hospital, Boston, MA, US

4, Department of Psychiatry and Psychotherapy, Universitätsmedizin Berlin Campus Charité Mitte, Berlin, DE

5, Medical and Population Genetics, Broad Institute, Cambridge, MA, US

6, Department of Psychiatry, Psychosomatics and Psychotherapy, University of Wurzburg, Wurzburg, DE

7, Centre for Psychiatry Research, Department of Clinical Neuroscience, Karolinska Institutet, Stockholm, SE

8, Department of Biomedicine, Aarhus University, Aarhus, DK

9, Dept of Biological Psychology & EMGO+ Institute for Health and Care Research, Vrije Universiteit Amsterdam, Amsterdam, NL

10, Division of Psychiatry, University of Edinburgh, Edinburgh, GB

11, Centre for Integrated Register-based Research, Aarhus University, Aarhus, DK

12, National Centre for Register-Based Research, Aarhus University, Aarhus, DK

13, iPSYCH, The Lundbeck Foundation Initiative for Integrative Psychiatric Research,, DK

14, Discipline of Psychiatry, University of Adelaide, Adelaide, SA, AU

15, Department of Translational Research in Psychiatry, Max Planck Institute of Psychiatry, Munich, DE

16, Department of Neurology, Klinikum rechts der Isar, Technical University of Munich, Munich, DE

17, Department of Psychiatry, Virginia Commonwealth University, Richmond, VA, US

18, Center for Neonatal Screening, Department for Congenital Disorders, Statens Serum Institut, Copenhagen, DK

19, Department of Psychiatry, Vrije Universiteit Medical Center and GGZ inGeest, Amsterdam, NL

20, Virginia Institute for Psychiatric and Behavior Genetics, Richmond, VA, US

21, Department of Psychiatry and Behavioral Sciences, Emory University School of Medicine, Atlanta, GA, US

22, Department of Medical Epidemiology and Biostatistics, Karolinska Institutet, Stockholm, SE

23, Department of Clinical Medicine, Translational Neuropsychiatry Unit, Aarhus University, Aarhus, DK

24, iSEQ, Centre for Integrative Sequencing, Aarhus University, Aarhus, DK

25, Human Genetics, Wellcome Trust Sanger Institute, Cambridge, GB

26, Statistical genomics and systems genetics, European Bioinformatics Institute (EMBL-EBI), Cambridge, GB

27, Department of Psychiatry, University Hospital of Lausanne, Prilly, Vaud, CH

28, Social Genetic and Developmental Psychiatry Centre, King's College London, London, GB

29, Genetics and Computational Biology, QIMR Berghofer Medical Research Institute, Brisbane, QLD, AU

30, Centre for Advanced Imaging, The University of Queensland, Brisbane, QLD, AU

31, Psychological Medicine, Cardiff University, Cardiff, GB

32, Center for Genomic and Computational Biology, Duke University, Durham, NC, US

33, Department of Pediatrics, Division of Medical Genetics, Duke University, Durham, NC, US

34, Centre for Cognitive Ageing and Cognitive Epidemiology, University of Edinburgh, Edinburgh, GB

35, Institute of Human Genetics, University of Bonn, School of Medicine & University Hospital Bonn, Bonn, DE

36, Epidemiology, Erasmus MC, Rotterdam, Zuid-Holland, NL

37, Psychiatry, Dokuz Eylul University School Of Medicine, Izmir, TR

38, Department of Psychiatry, Massachusetts General Hospital, Boston, MA, US

39, Psychiatric and Neurodevelopmental Genetics Unit (PNGU), Massachusetts General Hospital, Boston, MA, US

40, Stanley Center for Psychiatric Research, Broad Institute, Cambridge, MA, US

41, Neuroscience and Mental Health, Cardiff University, Cardiff, GB

42, Bioinformatics, University of British Columbia, Vancouver, BC, CA

43, Department of Epidemiology, Harvard T.H. Chan School of Public Health, Boston, MA, US

44, Department of Mathematics, Massachusetts Institute of Technology, Cambridge, MA, US

45, Department of Genetic Epidemiology in Psychiatry, Central Institute of Mental Health, Medical Faculty Mannheim, Heidelberg University, Mannheim, Baden-Württemberg, DE

46, Department of Psychiatry (UPK), University of Basel, Basel, CH

47, Department of Biomedicine, University of Basel, Basel, CH

48, Centre for Human Genetics, University of Marburg, Marburg, DE

49, Department of Psychiatry, Trinity College Dublin, Dublin, IE

50, Psychiatry & Behavioral Sciences, Johns Hopkins University, Baltimore, MD, US

51, Bioinformatics Research Centre, Aarhus University, Aarhus, DK

52, Institute of Genetic Medicine, Newcastle University, Newcastle upon Tyne, GB

53, Danish Headache Centre, Department of Neurology, Rigshospitalet, Glostrup, DK

54, Institute of Biological Psychiatry, Mental Health Center Sct. Hans, Mental Health Services Capital Region of Denmark, Copenhagen, DK

55, iPSYCH, The Lundbeck Foundation Initiative for Psychiatric Research, Copenhagen, DK

56, Brain and Mind Centre, University of Sydney, Sydney, NSW, AU

57, Interfaculty Institute for Genetics and Functional Genomics, Department of Functional Genomics, University Medicine and Ernst Moritz Arndt University Greifswald, Greifswald, Mecklenburg-Vorpommern, DE

58, Roche Pharmaceutical Research and Early Development, Pharmaceutical Sciences, Roche Innovation Center Basel, F. Hoffmann-La Roche Ltd, Basel, CH

59, Max Planck Institute of Psychiatry, Munich, DE

60, MRC Centre for Neuropsychiatric Genetics and Genomics, Cardiff University, Cardiff, GB

61, Department of Psychological Medicine, University of Worcester, Worcester, GB

62, Division of Research, Kaiser Permanente Northern California, Oakland, CA, US

63, Psychiatry & The Behavioral Sciences, University of Southern California, Los Angeles, CA, US

64, Department of Biomedical Informatics, Harvard Medical School, Boston, MA, US

65, Department of Medicine, Brigham and Women's Hospital, Boston, MA, US

66, Informatics Program, Boston Children's Hospital, Boston, MA, US

67, Wellcome Trust Centre for Human Genetics, University of Oxford, Oxford, GB

68, Institute of Social and Preventive Medicine (IUMSP), University Hospital of Lausanne, Lausanne, VD, CH

69, Swiss Institute of Bioinformatics, Lausanne, VD, CH

70, Division of Psychiatry, Centre for Clinical Brain Sciences, University of Edinburgh, Edinburgh, GB

71, Mental Health, NHS 24, Glasgow, GB

72, Department of Psychiatry and Psychotherapy, University of Bonn, Bonn, DE

73, Statistics, University of Oxford, Oxford, GB

74, Psychiatry, Columbia University College of Physicians and Surgeons, New York, NY, US

75, School of Psychology and Counseling, Queensland University of Technology, Brisbane, QLD, AU

76, Child and Youth Mental Health Service, Children's Health Queensland Hospital and Health Service, South Brisbane, QLD, AU

77, Child Health Research Centre, University of Queensland, Brisbane, QLD, AU

78, Estonian Genome Center, University of Tartu, Tartu, EE

79, Medical Genetics, University of British Columbia, Vancouver, BC, CA

80, Statistics, University of British Columbia, Vancouver, BC, CA

81, DZHK (German Centre for Cardiovascular Research), Partner Site Greifswald, University Medicine, University Medicine Greifswald, Greifswald, Mecklenburg-Vorpommern, DE

82, Institute of Clinical Chemistry and Laboratory Medicine, University Medicine Greifswald, Greifswald, Mecklenburg-Vorpommern, DE

83, Institute of Health and Biomedical Innovation, Queensland University of Technology, Brisbane, QLD, AU

84, Humus, Reykjavik, IS

85, Virginia Institute for Psychiatric & Behavioral Genetics, Virginia Commonwealth University, Richmond, VA, US

86, Clinical Genetics, Vrije Universiteit Medical Center, Amsterdam, NL

87, Complex Trait Genetics, Vrije Universiteit Amsterdam, Amsterdam, NL

88, Solid Biosciences, Boston, MA, US

89, Department of Psychiatry, Washington University in Saint Louis School of Medicine, Saint Louis, MO, US

90, Department of Biochemistry and Molecular Biology II, Institute of Neurosciences, Center for Biomedical Research, University of Granada, Granada, ES

91, Department of Psychiatry, University of Groningen, University Medical Center Groningen, Groningen, NL

92, Department of Psychiatry and Psychotherapy, University Hospital, Ludwig Maximilian University Munich, Munich, DE

93, Institute of Psychiatric Phenomics and Genomics (IPPG), University Hospital, Ludwig Maximilian University Munich, Munich, DE

94, Division of Cancer Epidemiology and Genetics, National Cancer Institute, Bethesda, MD, US

95, Behavioral Health Services, Kaiser Permanente Washington, Seattle, WA, US

96, Faculty of Medicine, Department of Psychiatry, University of Iceland, Reykjavik, IS

97, School of Medicine and Dentistry, James Cook University, Townsville, QLD, AU

98, Institute of Health and Wellbeing, University of Glasgow, Glasgow, GB

99, deCODE Genetics / Amgen, Reykjavik, IS

100, College of Biomedical and Life Sciences, Cardiff University, Cardiff, GB

101, Institute of Epidemiology and Social Medicine, University of Münster, Münster, Nordrhein-Westfalen, DE

102, Institute for Community Medicine, University Medicine Greifswald, Greifswald, Mecklenburg-Vorpommern, DE

103, Department of Psychiatry, University of California, San Diego, San Diego, CA, US

104, KG Jebsen Centre for Psychosis Research, Norway Division of Mental Health and Addiction, Oslo University Hospital, Oslo, NO

105, Medical Genetics Section, CGEM, IGMM, University of Edinburgh, Edinburgh, GB

106, Clinical Neurosciences, University of Cambridge, Cambridge, GB

107, Internal Medicine, Erasmus MC, Rotterdam, Zuid-Holland, NL

108, Roche Pharmaceutical Research and Early Development, Neuroscience, Ophthalmology and Rare Diseases Discovery & Translational Medicine Area, Roche Innovation Center Basel, F. Hoffmann-La Roche Ltd, Basel, CH

109, Department of Psychiatry and Psychotherapy, University Medicine Greifswald, Greifswald, Mecklenburg-Vorpommern, DE

110, Department of Psychiatry, Leiden University Medical Center, Leiden, NL

111, Virginia Institute for Psychiatric & Behavioral Genetics, Virginia Commonwealth University, Richmond, VA, US

112, Computational Sciences Center of Emphasis, Pfizer Global Research and Development, Cambridge, MA, US

113, Institute for Molecular Bioscience; Queensland Brain Institute, The University of Queensland, Brisbane, QLD, AU

114, Department of Psychiatry, University of Münster, Münster, Nordrhein-Westfalen, DE

115, Department of Psychiatry, University of Münster, Münster, DE

116, Department of Psychiatry, Melbourne Medical School, University of Melbourne, Melbourne, AU

117, Florey Institute for Neuroscience and Mental Health, University of Melbourne, Melbourne, AU

118, Institute of Medical Genetics and Pathology, University Hospital Basel, University of Basel, Basel, CH

119, Institute of Neuroscience and Medicine (INM-1), Research Center Juelich, Juelich, DE

120, Amsterdam Public Health Institute, Vrije Universiteit Medical Center, Amsterdam, NL

121, Centre for Integrative Biology, Università degli Studi di Trento, Trento, Trentino-Alto Adige, IT

122, Department of Psychiatry and Psychotherapy, Medical Center - University of Freiburg, Faculty of Medicine, University of Freiburg, Freiburg, DE

123, Center for NeuroModulation, Faculty of Medicine, University of Freiburg, Freiburg, DE

124, Psychiatry, Kaiser Permanente Northern California, San Francisco, CA, US

125, Medical Research Council Human Genetics Unit, Institute of Genetics and Molecular Medicine, University of Edinburgh, Edinburgh, GB

126, Department of Psychiatry, University of Toronto, Toronto, ON, CA

127, Centre for Addiction and Mental Health, Toronto, ON, CA

128, Division of Psychiatry, University College London, London, GB

129, Neuroscience Therapeutic Area, Janssen Research and Development, LLC, Titusville, NJ, US

130, Institute of Molecular and Cell Biology, University of Tartu, Tartu, EE

131, Psychosis Research Unit, Aarhus University Hospital, Risskov, Aarhus, DK

132, Munich Cluster for Systems Neurology (SyNergy), Munich, DE

133, University of Liverpool, Liverpool, GB

134, Mental Health Center Copenhagen, Copenhagen Universtity Hospital, Copenhagen, DK

135, Human Genetics and Computational Biomedicine, Pfizer Global Research and Development, Groton, CT, US

136, Psychiatry, Harvard Medical School, Boston, MA, US

137, Psychiatry, University of Iowa, Iowa City, IA, US

138, Department of Psychiatry and Behavioral Sciences, Johns Hopkins University, Baltimore, MD, US

139, Department of Psychiatry and Psychotherapy, University Medical Center Göttingen, Goettingen, Niedersachsen, DE

140, Human Genetics Branch, NIMH Division of Intramural Research Programs, Bethesda, MD, US

141, Faculty of Medicine, University of Iceland, Reykjavik, IS

142, Child and Adolescent Psychiatry, Erasmus MC, Rotterdam, Zuid-Holland, NL

143, Psychiatry, Erasmus MC, Rotterdam, Zuid-Holland, NL

144, Psychiatry, Dalhousie University, Halifax, NS, CA

145, Division of Epidemiology, New York State Psychiatric Institute, New York, NY, US

146, Department of Clinical Medicine, University of Copenhagen, Copenhagen, DK

147, Department of Medical & Molecular Genetics, King's College London, London, GB

148, Psychiatry & Behavioral Sciences, Stanford University, Stanford, CA, US

149, NIHR Maudsley Biomedical Research Centre, King's College London, London, GB

150, Genetics, University of North Carolina at Chapel Hill, Chapel Hill, NC, US

151, Psychiatry, University of North Carolina at Chapel Hill, Chapel Hill, NC, US

**Supplementary Note**

*Defining reported trauma exposure*

In this study, we sought to define reported trauma exposure with the aim of obtaining a single binary variable for stratification that reflected an overall exposure to a severe and potentially depressogenic environment. In doing so, we consulted a number of experts in the field of early-life trauma and stressful life events (including authors AD, BM, and MH among others). The complexity of defining reported trauma exposure is apparent from the multiple approaches that have been used in the field, and we do not suggest that the definition used here represents a gold-standard. However, we believe this is a reasonable definition of reported exposure given the necessary limitations of obtaining such data in a biobank-scale dataset.

Our definition included only items that were enriched in cases in this cohort, which we defined as having an OR > 2.5. This threshold results in only considering types of trauma more common in major depressive disorder (MDD) cases than in controls (Supplementary Table 2b). A side-effect of this is that the items on which we focus are more enriched in females than in males. As such, our enrichment for reported trauma exposure also reflects an enrichment for female sex, and for types of trauma that are more commonly reported by women.

We then defined individuals reporting two types of traumatic experience as reporters and those reporting no traumatic experiences as non-reporters. Individuals reporting only a single type of trauma were excluded. We did this because we wished to capture severe trauma, and felt that a single type of exposure may not represent such. This is imperfect, for a number of reasons. A single type of trauma may be depressogenic for a given individual, regardless of its wider effect in the population; however, the false positive rate of this more relaxed definition will be higher than a multi-item measure, given that 50% of the sample reported at least one traumatic experience. Reporting a type of trauma does not capture the severity of the trauma; however, the data available in the UK Biobank does not allow for a robust examination of trauma severity, and so multiple trauma reports are the best proxy available.

We included three measures of sexual trauma in our trauma definition. These are correlated, and so some participants may be defined as reporting trauma exposure from a single incident. However, the different sexual traumas are not nested; for example, within the analysed sample, only 3,758/7,179 (52%) individuals endorsing "interference by a partner" also endorsed "victim of sexual assault". This may result from semantic and contextual differences in how the questions were asked. The full sexual assault question is "*In your life, have you been a victim of a sexual assault, whether by a stranger or someone you knew"*, and the sexual interference question is "*Since I was sixteen, a partner or ex-partner sexually interfered with me, or forced me to have sex against my wishes"*. It is possible that participants felt that the first question encompassed more behaviours than the second. As such, we considered including all three items was justified, as excluding (for example) "interference by a partner" because it was correlated with "victim of sexual assault" could result in misclassification of individuals reporting the former but not the latter.

*Overlap with PGC major depression GWAS*

The PGC major depression GWAS contained participants from UK Biobank ^1^. Overlap of this kind upwardly biases results from polygenic risk scoring. Accordingly, we used a restricted set of summary statistics without these individuals (but including individuals from 23andMe ^2^). This addressed the major source of overlap between these cohorts. Further overlap may exist, but is difficult to quantify, as individual-level data is not available for all individuals in the PGC major depression GWAS. However, the deviation of the LD Score genetic covariance intercept from 0 represents a crude measure of sample overlap, assuming there is no shared confounding (such as population stratification) between the cohorts tested ^3,4^. The genetic covariance intercept in this case was -0.0061 (SE: 0.0063), indicating any remaining sample overlap is negligible.

*Additive interactions - linear regression or RERI?*

Linear regression was used in this analysis to test interaction as deviation from additivity, as has been used previously ^5^. However, an alternative method would be to calculate relative excess risk from interactions (RERI; ^6^), as has also been used previously ^7^. The use of linear regression in this context is less well-described than RERI, and may give an inaccurate estimate of interaction when there are sizable differences between the proportion of cases in the sample and in the population. However, there is only a minor difference between the proportion of cases in the sample (31.7%) and the population incidence of self-reported depression in England (28% ^8^), so this limitation is unlikely to affect our additive interaction analysis. As such, both methods were used.

**Supplementary Methods**

*Further information on main analyses*

*Phenotype distribution*

Participants were compared across a number of standard demographic variables and common correlates of MDD: sex, age (at questionnaire), education (university degree vs. not), neighbourhood socioeconomic status (SES, as Townsend deprivation index ^9^) and BMI (recorded from measurements taken at the initial recruitment of the participants into the biobank). For dichotomous variables (sex and education), comparisons were made using chi-square tests. For approximately continuous variables (age, SES and BMI), the skewness and kurtosis of the distribution was checked, and roughly normal variables (absolute values of skewness (as b_1_) and kurtosis (as b_2_) <= 2; ^10^) were compared using Welch's t-tests. Non-normal continuous variables were compared using Mann-Whitney U tests. All comparisons were performed in R.3.4.1, using skewness and kurtosis calculations from the e1071 package ^10–12^. An additional breakdown of individual trauma items by sex was performed.

*Genome Wide Association Studies (GWAS)*

GWAS were performed using linear regressions on imputed genotype dosages in BGenie v1.2 ^13^, with residualised phenotypes. Deviance residuals were obtained from logistic regressions in R.3.4.1 ^11^. Six principal components were used because investigations suggested this was necessary to control for geographical variation in the dataset, and that increasing this number had negligible additional benefits (data not shown).

Results from each GWAS were clumped to define genetic loci in PLINK2 ^14^. Loci were defined following established protocols ^1^. Each locus comprised all variants with p < 0.0001 in linkage disequilibrium (r^2^ > 0.1 in European subjects from the 1000 Genomes Phase 3 release ^15^) with a nearby (< 3Mb) variant with a lower p-value. Neighbouring (< 50kb) or overlapping clumps were merged using bedtools ^16^. Loci were annotated using RegionAnnotator v1.63 (<https://github.com/ivankosmos/RegionAnnotator>) to identify proximal (< 100kb from loci boundaries) features of interest using data from the NHGRI-EBI GWAS Catalog; OMIM; GENCODE genes; genes previously implicated in autism and/or intellectual disability; copy-number variants previously implicated in psychiatric disorders; and mouse knockout phenotypes.

*Sensitivity analyses*

Sensitivity analyses were performed as described below. For all analyses, phenotypes were residualised using the same process and covariates as described in the main text.

*Reported trauma exposure*

Three sets of analyses were performed comparing (i) all individuals reporting trauma exposure with those not reporting exposure, (ii) limiting just to MDD cases, and (iii) limiting just to controls. Participants included in the overall reported trauma exposure analysis, and the analyses stratified by MDD, were compared across the same phenotypic variables used in the main analysis. All analyses performed in the main paper were repeated focussed on reported trauma exposure, with the exception of stratified heritability analyses, as we felt the results of such analyses would be difficult to interpret.

*Covarying for age, education, neighbourhood SES and BMI*

Analyses of MDD (overall, in individuals reporting trauma exposure, in individuals not reporting trauma exposure) and reported trauma exposure (overall, in cases, in controls) were repeated, retaining all previous covariates and including as further covariates age (at questionnaire), neighbourhood socioeconomic status (SES, as Townsend deprivation index ^9^), BMI (at baseline assessment), and a binary variable of education (university degree vs. not). All analyses performed in the main paper (and as described above, focussed on reported trauma exposure) were repeated.

*Downsampled cohorts*

Most of the sample with data both on MDD symptoms and on reported trauma status were controls who did not report trauma (Table 1). To assess whether this disbalance in sample status affected our results, analyses of genetic correlations between external phenotypes and MDD (i) overall, (ii) in individuals reporting trauma exposure, and (iii) in individuals not reporting trauma exposure were repeated using ten downsampled cohorts. Downsampled cohorts were produced by downsampling both MDD case groups (reporting and not reporting trauma exposure) and the control group not reporting trauma exposure to 9,487 (the smallest group, controls reporting trauma exposure). Downsampling was performed in R 3.4.1, using random selections of individuals from each group, and was repeated ten times to reduce selection biases ^11^. Genetic correlations with all external phenotypes were performed in LD Score for each of the six analyses (MDD overall and stratified, and reported trauma exposure overall and stratified) as detailed in the main paper. The mean average SNP-heritability estimates and mean average standard errors from each of the ten cohorts were calculated. From each analysis, the average genetic correlation with each external phenotype was compared to the relevant correlation from the main analyses ^17^. The difference between correlations with MDD in individuals reporting and not reporting trauma exposure were also compared, and then these differences were compared to those from the main analysis. All comparisons used two-sample z-tests. Use of the block jackknife was not possible as the results were averaged across downsampled cohorts. Equivalent comparisons were made for analyses of reported trauma exposure in cases and in controls.

*Alternative definitions of reported trauma exposure*

In order to test the robustness of our main finding (that the SNP-heritability of MDD is greater in individuals reporting trauma exposure compared to those who do not), we repeated this analysis with three alternative definitions of reported trauma exposure. Our first two alternatives altered the threshold for including MDD-relevant traumas, firstly decreasing the threshold from reporting two such traumas to reporting one, and secondly increasing it to reporting three MDD-relevant traumas. Our third alternative altered the definition itself to focus only on childhood traumas. We considered all five childhood traumas, and defined reported trauma exposure as a report of any of these five (Supplementary Table 2a). Traumas in adulthood and PTSD-relevant traumas were not considered for the purpose of this final alternative (so individuals defined as not reporting trauma on this measure may still report traumatic experiences later in life).

*Heritability analyses stratified by reported trauma exposure*

We aimed to compare the SNP-heritability of MDD in individuals reporting trauma exposure with those not doing so. Typically, such a comparison could be achieved by converting the SNP-heritability estimates from the observed scale (which is dependent on the proportion of cases in the cohort) to the liability scale (independent of the proportion of cases in the study and of the population prevalence). We calculated the proportion of individuals with MDD in each group (i.e. we converted Table 1 from counts to proportions). We then assumed that the population prevalence of self-reported MDD = 28% ^8^ and that individuals with trauma exposure were sampled representatively from the population in cases and in controls. This allows the calculation of population prevalences for MDD in individuals reporting (52%) and not reporting trauma exposure (17%). With these estimates, and observed scale SNP-heritability estimates from BOLT-LMM, we estimated of liability scale SNP-heritability estimates for MDD in individuals reporting and not reporting trauma exposure, and compared them using a two-sample z-test (Supplementary Table 7).

However, the conversion from the observed to the liability scale assumes the genetic and environmental influences on the trait are independent - the correlation between MDD and reported trauma exposure violates this underlying assumption ^18–20^. Furthermore, the sample is not divided by an purely environmental trait, because trauma exposure is itself heritable. The impact of these concerns on estimates of the SNP heritability in each group is not intuitive.

Therefore, we performed a simulation study of SNP-level data in line with previous work ^21^. In this simulation, we assumed the phenotypic link between MDD and reported trauma exposure was attributable to sharing of genetic and environmental effects without interaction (that is, gene-environment correlation alone). The prevalence of MDD was set at $K_{MDD}=0.28$. The remaining simulation parameters were aligned with the empirical observations from the study: the prevalence of reported trauma exposure was set at $K_{T}=0.32$ (assuming accurate sampling of reported trauma exposure from the population of cases and of controls), while the liability-scale heritabilities were set at $h_{l,MDD}^{2}=0.20$ and $h_{l,T}^{2}=0.24$. BOLT-LMM was used to empirically estimate the genetic correlation between MDD and reported trauma exposure at $r_{g}=0.76$ and the environmental correlation at $r_{e}=0.31$. These values were used as the first parameterization (parametrization #1) and fixed the phenotypic OR between MDD and reported trauma exposure at 3.1. The empirical OR, however, was estimated at 5.2. Therefore, a second parameterization was simulated by increasing $r_{e}=0.5$ and thereby fixing the phenotypic OR at 5.2 (parametrization #2). For these parameterizations, 1,000 SNPs were simulated with random minor allele frequencies (MAF) uniformly distributed between 0.05 and 0.5. The SNP-effects for MDD ($\beta_{MDD}$) and reported trauma exposure ($\beta_{T}$) were drawn from a bivariate normal distribution with variances $h_{l,MDD}^{2}=0.20$ and $h_{l,T}^{2}=0.24$ and covariance$r_{g}\sqrt{h_{l,MDD}^{2}h_{l,T}^{2}}$. An individual was simulated by randomly assigning genotypes with probabilities in line with the MAFs. The genetic value for MDD was estimated as $g_{MDD}=\sum_{i} \beta_{MDD,i}({genotype}_{i}-2{MAF}_{i})/(2{MAF}_{i}(1-{MAF}_{i}))$ with $i$ iterating over the 1,000 SNPs, and the genetic values for reported trauma exposure were estimated analogously. The environmental values for MDD ($e_{MDD}$) and reported trauma exposure ($e_{T}$) were drawn from a bivariate normal distribution with variances $1-h_{l,MDD}^{2}$ and $1-h_{l,T}^{2}$ and covariance$r_{e}\sqrt{(1-h_{l,MDD}^{2})({1-h}_{l,T}^{2})}$ . The values on the liability scale were subsequently estimated as $l_{MDD}=g_{MDD}+e_{MDD}$ and $l_{T}=g_{T}+e_{T}$. MDD was set at 1 when $l_{MDD}\geq\phi^{-1}(1-K_{MDD})$ and 0 otherwise, while reported trauma exposure was set 1 when $l_{T}\geq\phi^{-1}(1-K_{T})$ and 0 otherwise ($\phi^{-1}$ is the standard normal cumulative distribution function). For both parametrizations, individuals were simulated one-by-one until we had collected 4,000 of each of: cases reporting trauma exposure (MDD_1_T_1_), controls reporting trauma exposure (MDD_0_T_1_), cases not reporting trauma exposure (MDD_1_T_0_), and controls not reporting trauma exposure (MDD_0_T_0_). The data were merged for MDD_1_T_1_-MDD_0_T_1_ and MDD_1_T_0_-MDD_0_T_0_. Subsequently, cross-product Haseman Elston regression was applied to estimate the heritability of MDD in individuals reporting trauma exposure and in individuals not reporting trauma exposure respectively. The observed scale heritabilities were converted to the liability scale based on a prevalence of MDD in individuals reporting trauma exposure of $K_{MDD|T=1}=0.45$ and in individuals not reporting trauma exposure of $K_{MDD|T=0}=0.20$, as follows from $K_{MDD}=0.28$ and $K_{T}=0.32$ with $OR=3.1$ (i.e. parametrization #1) ^22^. For parametrization #2, these respective prevalences were $K_{MDD|T=1}=0.52$ and $K_{MDD|T=0}=0.17$ (as follows from $K_{MDD}=0.28$ and $K_{T}=0.32$ with $OR=5.2$). For both parameterizations, simulations were repeated 100 times.

With these simulations an average liability scale heritability of MDD $\hat{h}_{l,MDD|T=1}^{2}=0.148$ (SE over 100 iterations of 0.002) was found in individuals reporting trauma exposure, and $\hat{h}_{l,MDD|T=0}^{2}=0.157$in individuals not reporting trauma exposure for parameterization #1. For parameterization #2, heritabilities were estimated as $\hat{h}_{l,MDD|T=1}^{2}=0.145 (0.002)$ and $\hat{h}_{l,MDD|T=0}^{2}=0.154 (0.001)$ respectively. Thus, the heritability estimates were slightly larger in individuals not reporting trauma exposure compared to individuals reporting trauma exposure. This contrasts our empirical findings where the heritability was larger in individuals reporting trauma exposure compared to individuals not reporting trauma exposure. Thus, these simulations suggest that our empirical findings were not directly attributable to bias from the heritable component of reported trauma exposure, nor by the transformation of the observed scale heritability to the liability scale. At the same time, however, we would like to emphasize that we did not address different sources of potential bias from other genetic architectures than those simulated, from intrinsic challenges of heritability estimation from case-control data ^21,23^, or from potential collider bias resulting from selection bias ^24^.

*Comparing two genetic correlations using block jackknife and LD Score*

Define four phenotypes: A, B, C, and D. We wished to compare the genetic correlation of A and B to the genetic correlation of C and D. Global estimates of these correlations, denoted $r\left( A,B \right)$ and $r\left( C,D \right)$, can be computed using LD Score. The same software can output jackknife delete values for genetic covariance: $G\left( A,B \right)$, $G\left( C,D \right)$, as well as for heritability: $H\left( A,B \right)$ and $H\left( C,D \right)$. These jackknife delete values are estimated by excluding blocks of values (here, number of blocks n = 200). The n-dimensional vectors $G\left( A,B \right)$, $G\left( C,D \right)$, $H\left( A,B \right)$ and $H\left( C,D \right)$ can be used to generate genetic correlation delete values $R\left( A,B \right)$ and $R\left( C,D \right)$. The difference between the global estimates $r\left( A,B \right)$ and $r\left( C,D \right)$ is $d\left( AB,CD \right)$ and the difference between the vectors $R\left( A,B \right)$ and $R\left( C,D \right)$ is $D\left( AB,CD \right)$. The global genetic correlation difference $d\left( AB,CD \right)$ and the delete values $D\left( AB,CD \right)$ are used to compute jackknife pseudovalues. The ith pseudovalue is:

$$P_{i}\left( AB,CD \right)=n\times d\left( AB,CD \right)-\left( n-1 \right)*D_{i}\left( AB,CD \right)$$

The mean and variance of the jackknife pseudovalues are:

$$m(AB,CD)=\frac{1}{n}\sum_{i=1}^{n} P_{i}(AB,CD)$$

$$v(AB,CD)=\frac{1}{n-1}\sum_{i=1}^{n} {(P_{i}(AB,CD)-m(AB,CD))}^{2}$$

The jackknife estimate of the difference between the two correlations m(AB,CD) can then be compared to test H0 : θ = θ_0_ (where θ_0_ = 0 for no difference between genetic correlations), and a p-value can be derived from the z statistic:

$$z(AB,CD)=\frac{m(AB,CD)-\theta_{0}}{\sqrt{(1/n)\times v(AB,CD)}}$$

*Gene environment interaction analyses (variant level)*

In addition to the analyses described in the main text, we performed exploratory analyses to determine if the variants with the strongest main effects in the analyses of MDD and reported trauma exposure (overall and stratified) showed interaction effects. Specifically, index SNPs with a nominally significant main effect (p<0.0001) in one or more of the six analyses were tested for SNP-by-reported trauma exposure interactions associated with MDD. (Similar analyses were performed assessing SNP-by-MDD interactions and reported trauma exposure, but results were effectively identical to those for SNP-by-reported trauma interactions and are not shown.) Interaction models were calculated in R.3.4.1 ^11^. SNP dosages were extracted from imputed data using PLINK2.0 ^14^. SNP, reported trauma and genetic principal components were mean-centred and scaled to have a standard deviation of 1 (i.e. converted to Z scores). To assess interactions as departure from multiplicativity, logistic regressions were performed regressing MDD on the main effect of reported trauma exposure, SNP, covariates (as used in the GWAS), SNP-by-reported trauma exposure interaction terms, SNP-by-covariate interaction terms and reported trauma exposure-by-covariate interaction terms ^25,26^. Further analyses were performed to assess interactions as departure from additivity as above using linear regressions. SNP-trauma interaction terms were considered experiment-wide significant if they passed Bonferroni correction for the 1,652 SNPs assessed (p ≈ 3x10^-5^), and genome-wide significant if p ≤ 5x10^-8^.

**Supplementary Results**

*Sensitivity analyses*

*Analyses focussed on reported trauma exposure*

Individuals reporting trauma exposure differed significantly from those who did not: they were mostly females, significantly younger, more likely to have a degree, came from more deprived neighbourhoods, and had higher BMI at recruitment (all p < 0.05; Supplementary Table 4). Genome-wide association analyses identified six significant loci when comparing individuals reporting trauma exposure to those who did not, which remained significant when assessed with logistic regression (Supplementary Figures 4-6). These loci have previously been implicated in genetic studies of a variety of phenotypes, including attention deficit disorder, schizophrenia and educational attainment, but did not overlap with the locus identified in the MDD analyses (Supplementary Table 8). No analysis showed evidence of genome-wide inflation that was attributable to confounding (95% confidence intervals of all regression intercepts from LD Score heritability estimation overlapped 1; Supplementary Table 7). The liability-scale SNP-heritability estimate for reported trauma exposure (24% [22-26%], assuming a population prevalence of 32%, equivalent to representative sampling from the population in cases and in controls) was in excess of that for MDD (20%, Z-test p = 0.006).

Genetic correlations were calculated between all internal phenotypes (Supplementary Table 10). The genetic correlation between reported trauma exposure in cases and in controls was high (r_g_ = 0.737 [95% CI: 0.493-0.981]; difference from 0: p = 3.33 x 10^-9^; difference from 1: p = 0.0349). Genetic correlations of reported trauma exposure with body composition phenotypes and with educational attainment were significantly larger in cases than the equivalent correlations in controls (Supplementary Figure 7; Supplementary Table 11).

Individuals with higher MDD PRS were more likely to report a trauma exposure, and a significant additive interaction term was observed from linear regression - the combined effect of PRS and reported trauma exposure was greater than the sum of the individual effects (beta > 0, Supplementary Table 12). However, the multiplicative interaction term was not significant (p > 0.01). Individuals with higher BMI risk scores were more likely to be report a trauma exposure, although this only passed correction for multiple testing in cases. Both the additive (beta > 0) and the multiplicative (OR > 0) interaction terms were significant, suggesting the combined risk of MDD from BMI PRS and reported trauma exposure together was greater than expected from both the sum of the individual risks and from their product, respectively (OR > 1).

*Covarying for BMI, age, SES and education*

Approximately 1% of the cohort did not provide information on one or more of these variables, resulting in a minor difference in sample size across all analyses (Supplementary Table 13).

No additional loci passed genome-wide significance in any analysis. The locus passing genome-wide significance in the overall MDD analysis, and four of the six loci from the overall reported trauma exposure analyses, remained significant when controlling for the additional covariates (Supplementary Table 14). There was no evidence of confounding introduced by controlling for these additional covariates (all LD Score intercepts contained 1; Supplementary Table 15).

Estimates of the SNP-heritability of MDD overall (19% [17-21%]) and in individuals reporting trauma exposure (22% [16-29%]) were attenuated compared to the overall analysis, but remained constant in those not reporting trauma exposure (12% [7-16%]; Supplementary Table 15). Despite this, the SNP-heritability of MDD remained significantly greater in individuals reporting trauma exposure than in those not reporting trauma exposure (p = 0.01). Similarly, estimates of the SNP-heritability of reported trauma exposure were attenuated (23% [21-25%]).

Genetic correlations with external phenotypes did not differ substantially with and without the additional covariates in any analysis (all z-test p < 0.05), with the exception of the correlations with BMI, body fat percentage and fat mass, all of which were significantly diminished in the overall MDD analysis when the additional covariates were included (z-test p = 0.01-0.03). The genetic correlation of waist circumference with MDD no longer differed significantly between individuals reporting and not reporting trauma exposure. In the analyses of reported trauma exposure, genetic correlations with BMI, fat mass, hip circumference and waist-hip ratio no longer differed significantly between cases and controls. However, the correlation between reported trauma exposure and total body (less head) bone mineral density became significantly larger in cases compared to controls (Supplementary Table 17).

PRS analyses differed only in analyses involving the BMI PRS. BMI PRS was no longer associated with MDD or reported trauma exposure in any analysis, and no interactions including the BMI PRS remained significant after correcting for multiple testing (Supplementary Table 18).

*Genetic correlations using downsampled cohorts*

Genetic correlation analyses with external phenotypes were rerun using ten cohorts downsampled such that each group had 9,487 participants (Supplementary Table 19). Mean average genetic correlations with MDD were attenuated across most phenotypes when compared to the original results, but these reductions were not significant in any instance (two-sample z-tests, all p > 0.05). Similarly, differences in correlations with MDD between individuals reporting trauma exposure and individuals not reporting trauma exposure were reduced compared to the original results, but not significantly so (two-sample z-tests, all p > 0.05). However, due to this reduction, no differences in genetic correlations with MDD between individuals reporting and not reporting trauma exposure remained significant after multiple testing correction in this downsampled cohort. As such, we conclude that the differences observed in the genetic correlations with MDD between individuals reporting and not reporting trauma exposure are robust, but their magnitude is likely to be increased by the differences in size between the different groups.

*SNP-heritability of MDD using alternative definitions of reported trauma exposure*

We repeated our analyses of SNP-heritability in MDD using alternative definitions of reported trauma exposure in order to test the robustness of this principal finding from the main paper. Results did not qualitatively differ from those in the main paper: for all definitions of reported trauma exposure, the SNP-heritability of MDD was significantly greater in individuals reporting trauma exposure than in those not doing so (Supplementary Table 20).

Altering the threshold for including MDD-relevant traumas resulted in a dosage-like effect, whereby increasing the number of reported traumas required increased the SNP-heritability of MDD in individuals reporting trauma exposure (21% to 24% to 28%). However, increasing the threshold reduced the number of individuals defined as reporting trauma exposure. Consequently, the power decreased with higher thresholds, as did the significance of the difference in MDD SNP-heritability between those reporting and not reporting trauma exposure.

Limiting the trauma items considered just to the childhood items altered the composition of both the group reporting and the group not reporting trauma exposure (Compare the two previous alternative definitions, in which the composition of the group not reporting trauma remained constant.) Although the difference in MDD SNP-heritability between the trauma-reporting and non-reporting groups was smaller using this final alternative definition, the greater power of this definition meant that the difference remained significant.

*SNP environment interaction analyses*
Analyses were performed for the 1,652 index SNPs (p < 0.0001 in at least one of the six analyses) to assess the association of SNP-trauma interactions with MDD. Multiplicative interaction effects at experiment-wide significance (p < 3x10^-5^) were observed at 78 variants, with the most significant interaction (with rs143276464) reaching genome-wide significance (p = 2.42x10^-9^; Supplementary Table 9). In the case of rs143276464, the probability of MDD increases with each additional minor allele in individuals reporting trauma exposure, but decreases with each additional minor allele in those who do not report exposure.

Additive interaction effects at experiment-wide significance (p < 3x10^-5^) were observed at 85 variants, although none reached genome-wide significance (Supplementary Table 9). 31/85 of the interactions with significant additive interaction effects also showed significant multiplicative interaction effects.

Replication of the observed interactions was sought in a previous analysis of genetic variant-by-trauma interactions with MDD (N = 3944, ^27^) and in similar data from Generation Scotland (N = 629, unpublished). No interaction was significantly associated in both replication datasets.
