## Supplementary Figures for "Genome-wide gene-environment analyses of major depressive disorder and reported lifetime traumatic experiences in UK Biobank"

Supplementary Figure 1

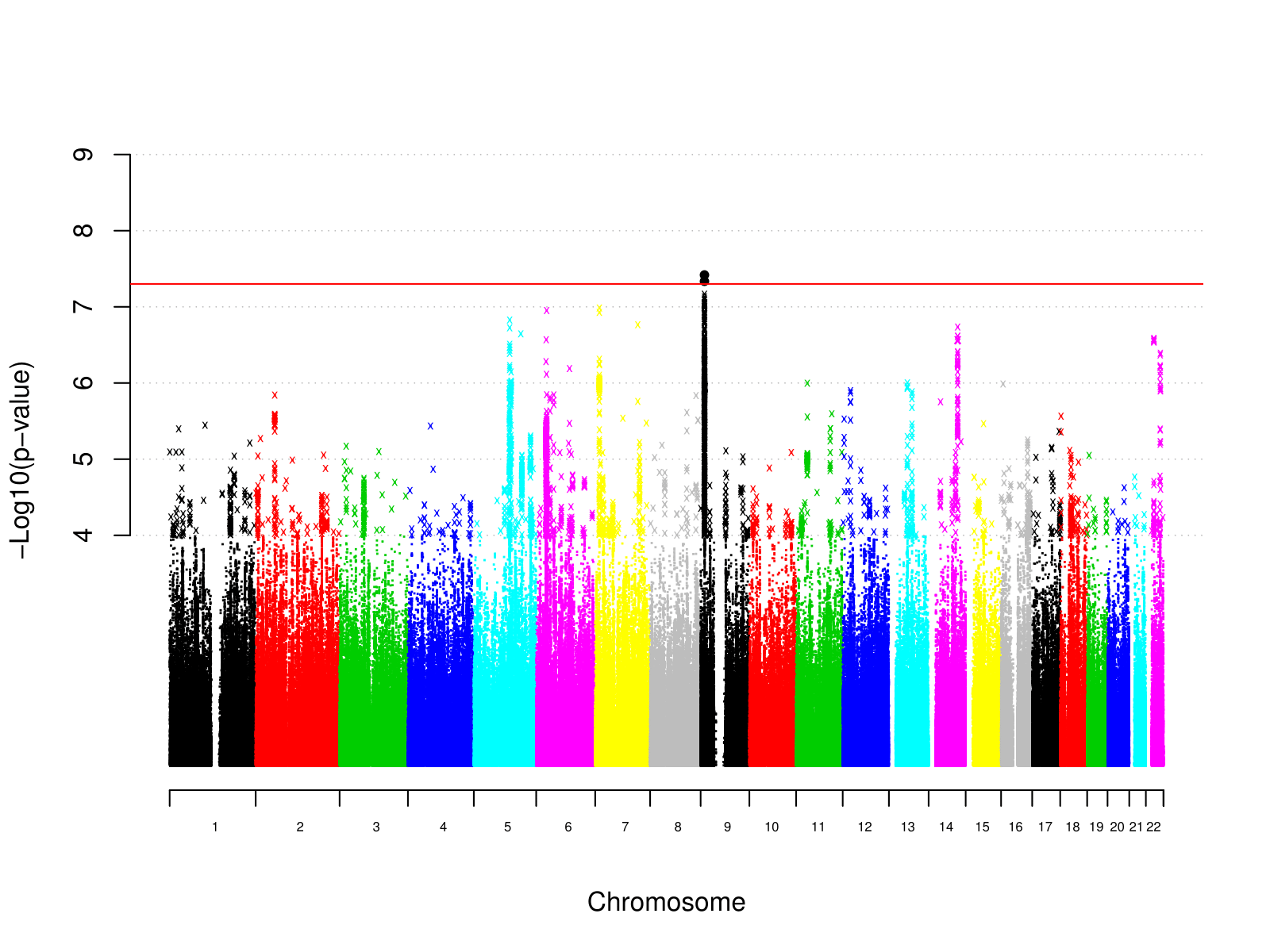

Supplementary Figure 1: Manhattan plot of individual variant GWAS results for MDD

Supplementary Figure 2

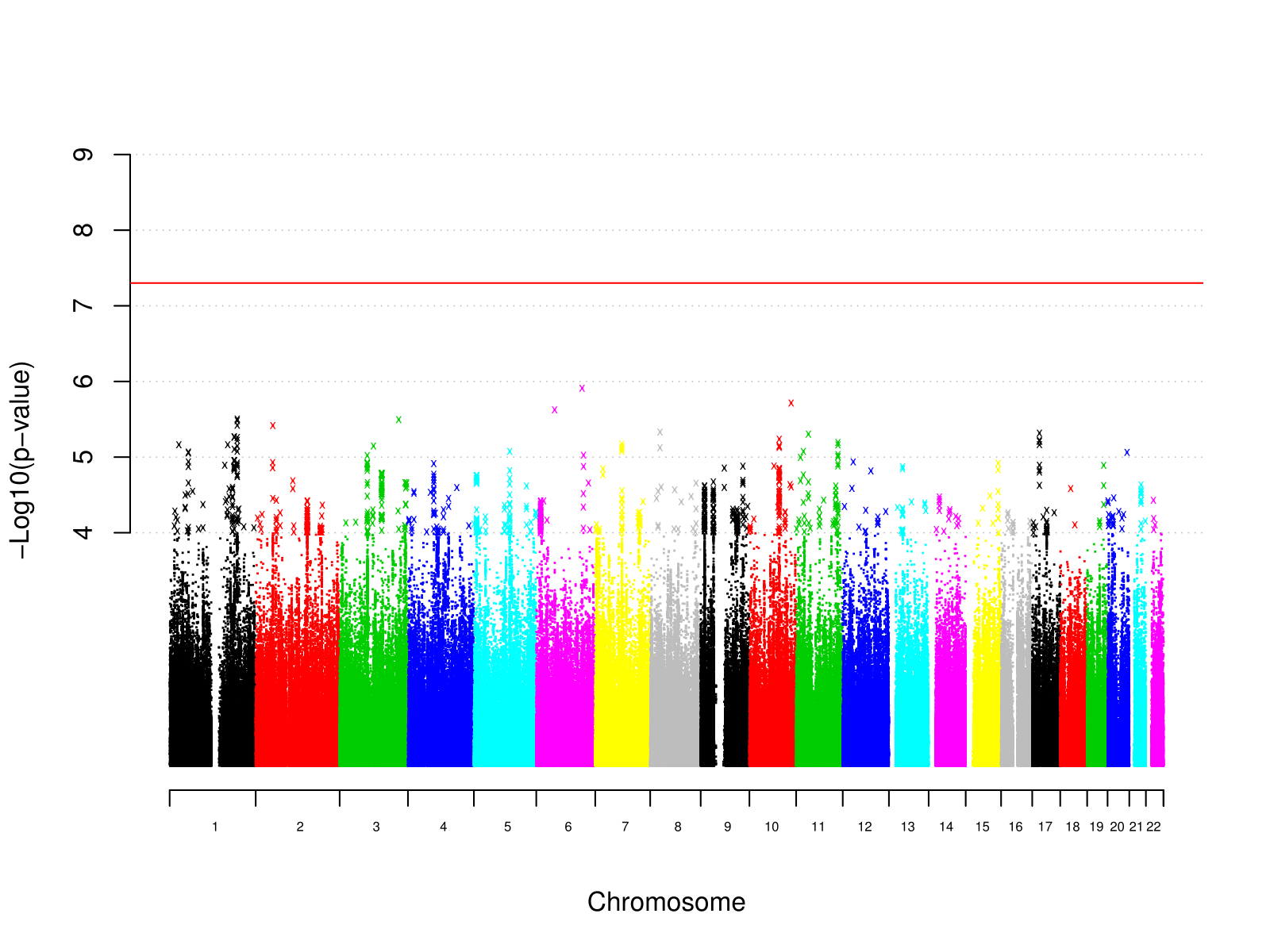

Supplementary Figure 2: Manhattan plot of individual variant GWAS results for MDD in individuals reporting trauma exposure

Supplementary Figure 3

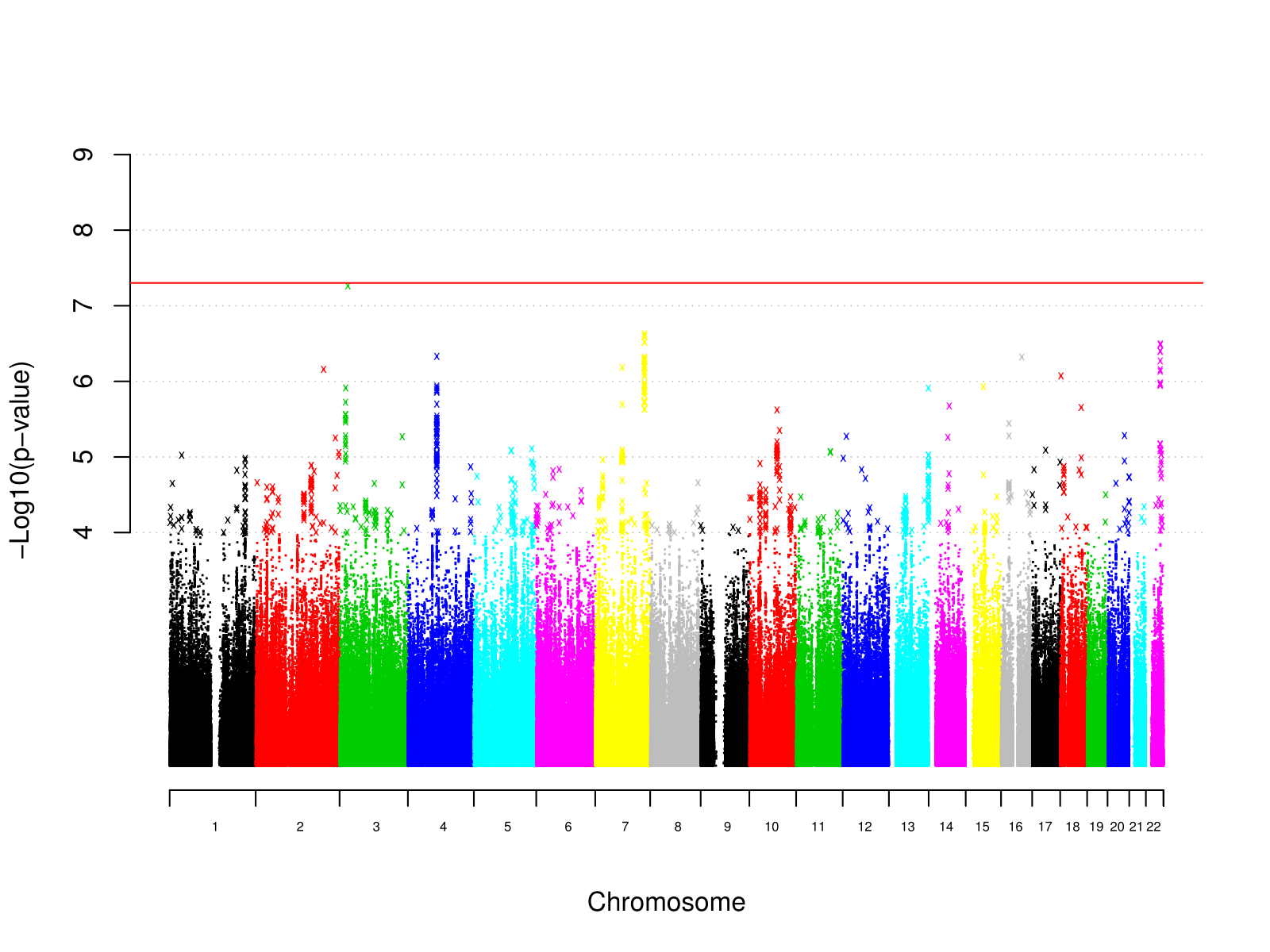

Supplementary Figure 3: Manhattan plot of individual variant GWAS results for MDD in individuals not reporting trauma exposure

Supplementary Figure 4

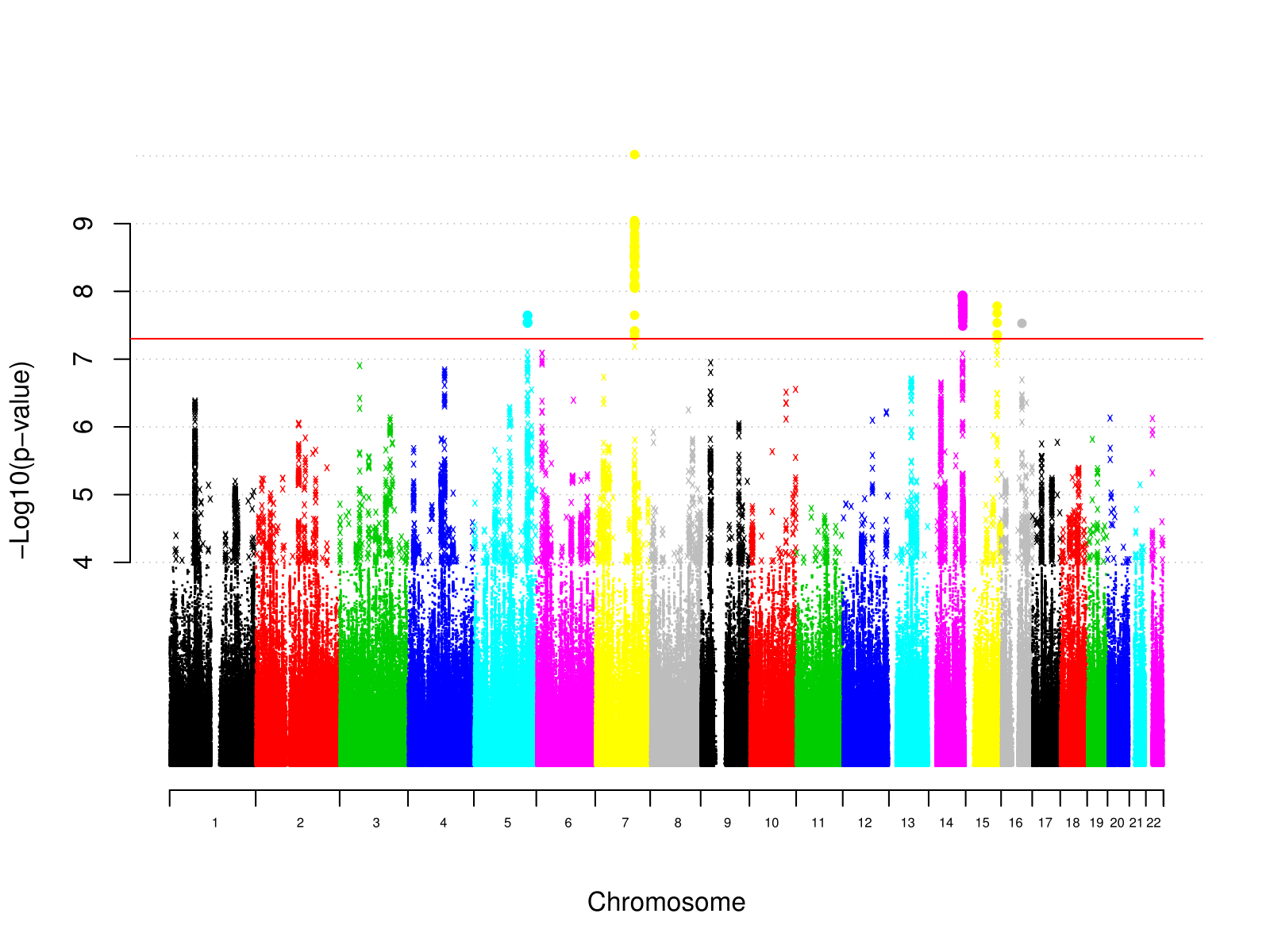

Supplementary Figure 4: Manhattan plot of individual variant GWAS results for reported trauma exposure

Supplementary Figure 5

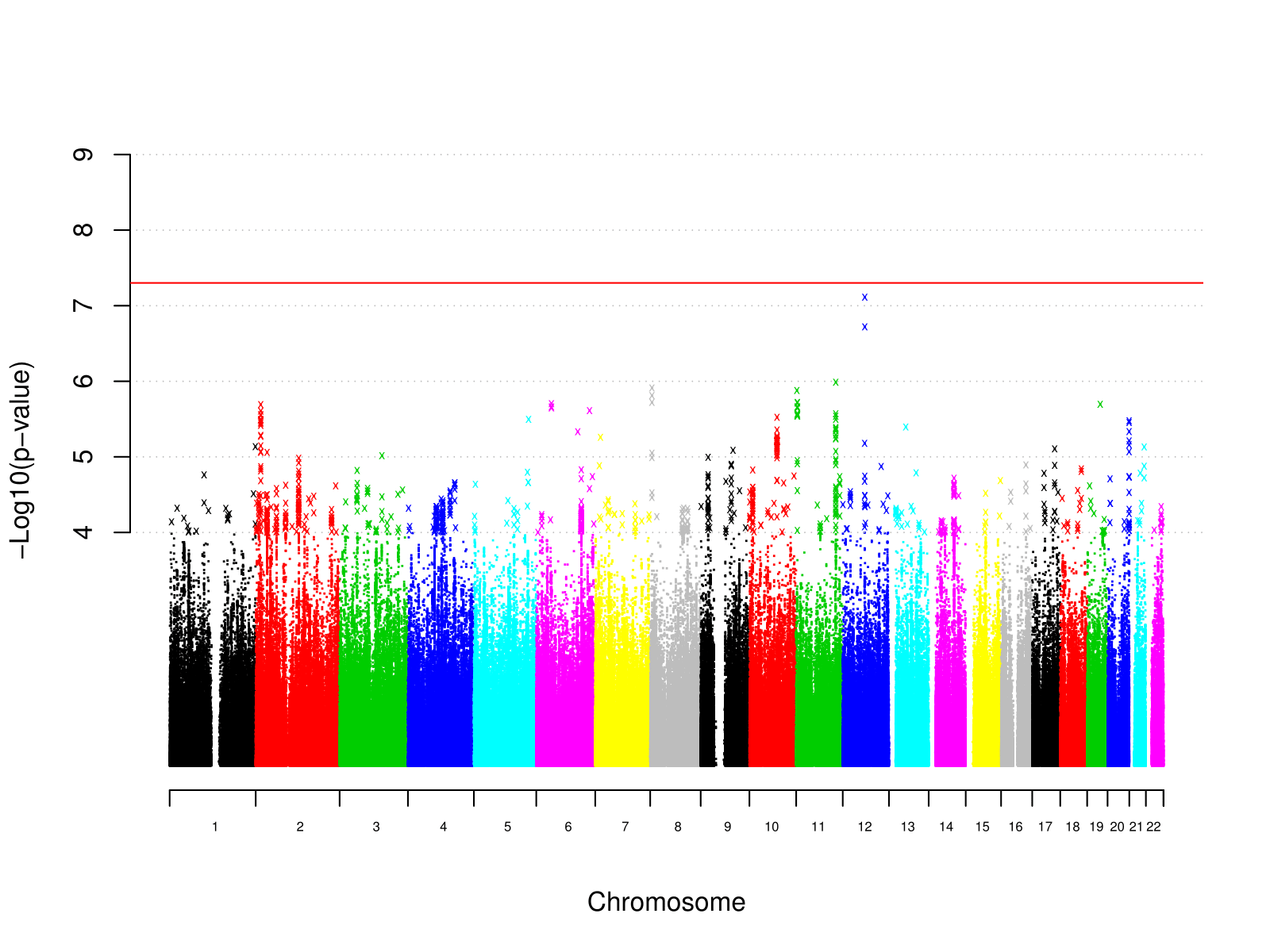

Supplementary Figure 5: Manhattan plot of individual variant GWAS results for reported trauma exposure in MDD cases

Supplementary Figure 6

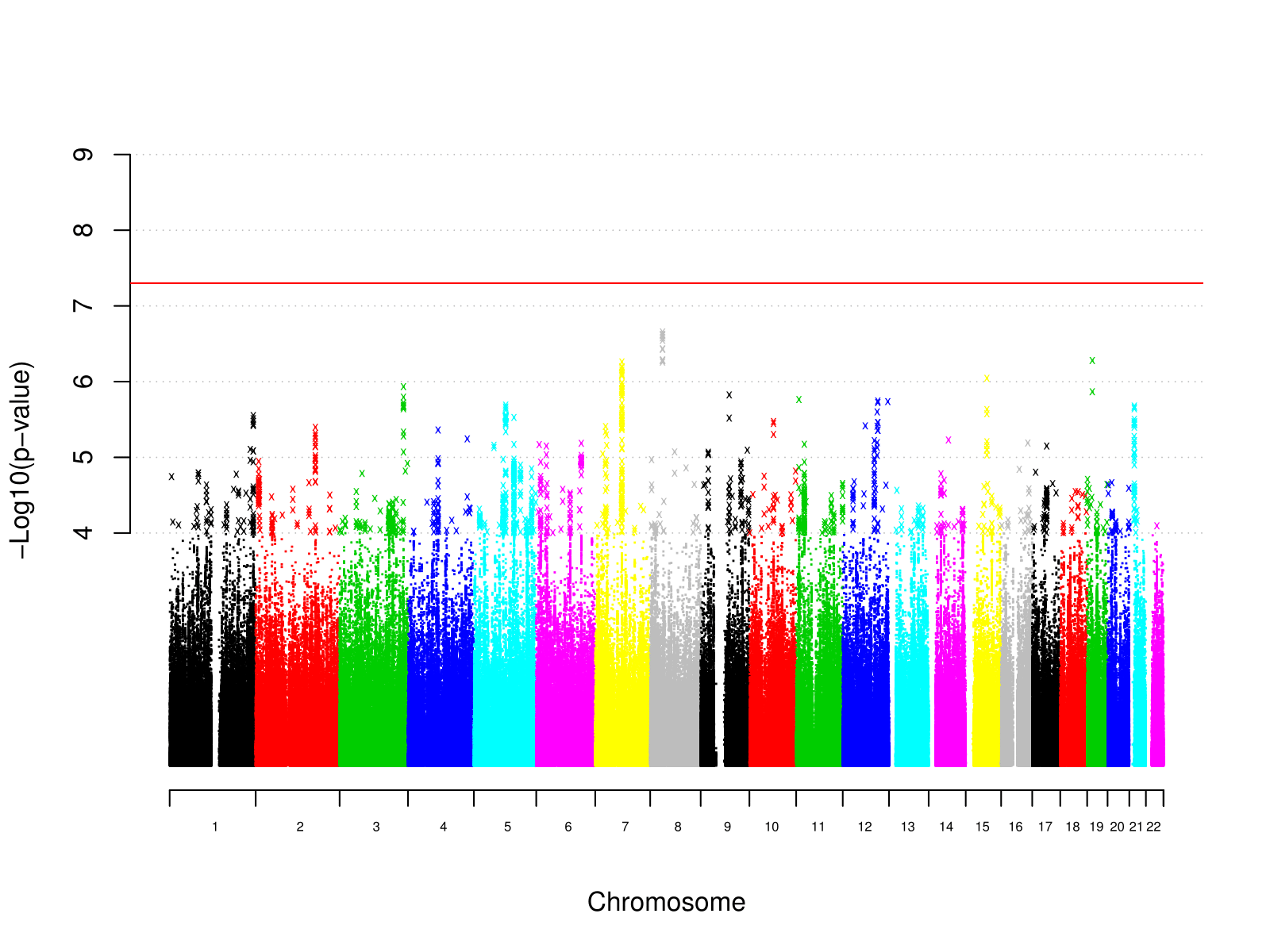

Supplementary Figure 6: Manhattan plot of individual variant GWAS results for reported trauma exposure in controls

Supplementary Figure 7

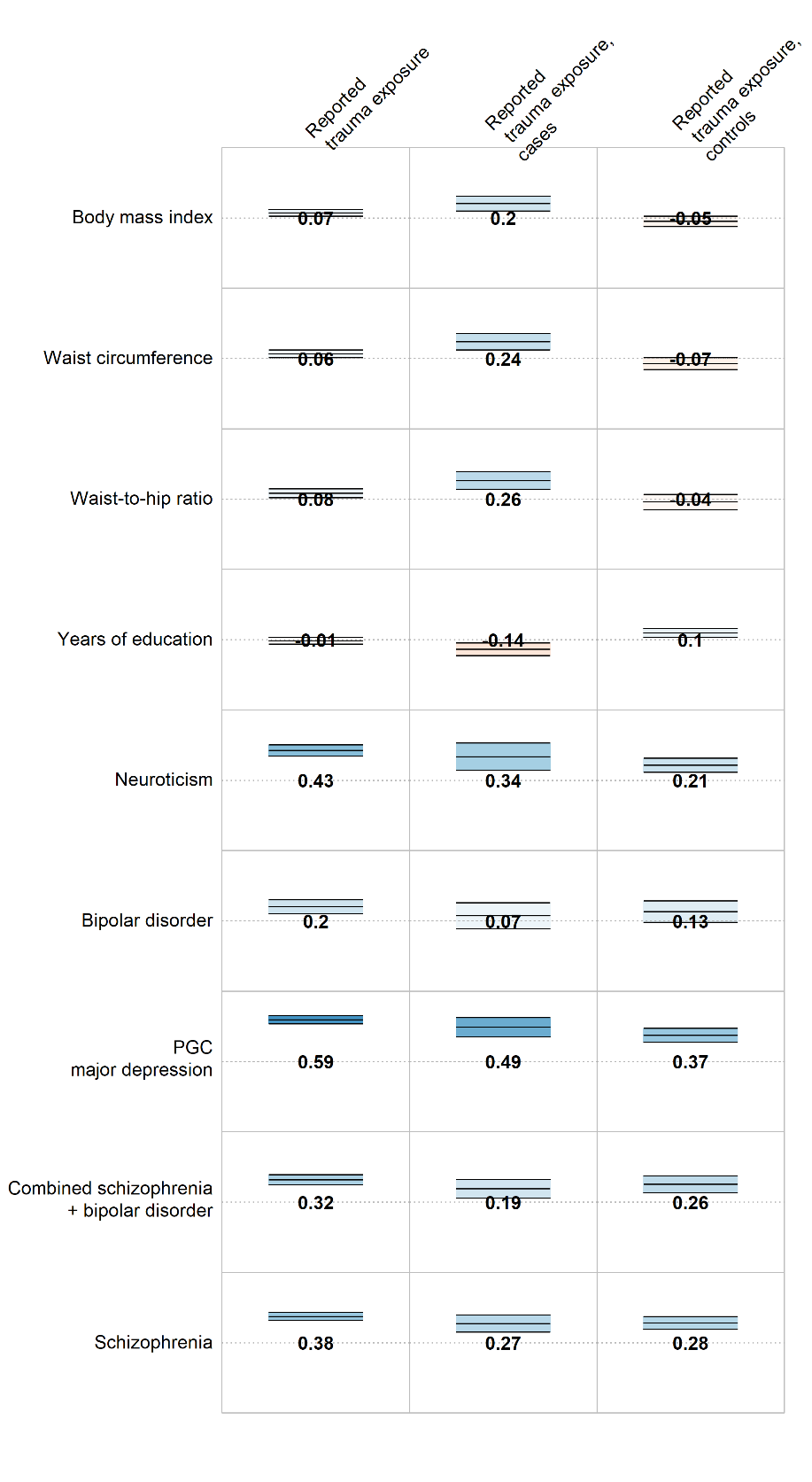

Supplementary Figure 7: Genetic correlations between external phenotypes and reported trauma exposure, differing by MDD case status (middle and right columns). Numbers = genetic correlations. Colour = direction of effect (blue = positive, red = negative). Colour intensity = size of correlation. Upper and lower bars are 95% confidence interval of genetic correlation.
